# A bacterial PrimPol-reverse transcriptase hybrid protein has a proofreading exonuclease activity that can be transferred to other reverse transcriptases

**DOI:** 10.1101/2025.09.07.674619

**Authors:** Seungjin Kim, Hengyi Xu, Georg Mohr, Jun Yao, Y. Jessie Zhang, Guzhen Cui, Alan M. Lambowitz

**Affiliations:** Department of Molecular Biosciences, University of Texas at Austin, Austin, TX 78712; School of Basic Medicine, Guizhou Medical University, Guiyang 561113, China; Department of Medicine / Oncology, University of Texas at Austin, Austin, TX 78712

**Keywords:** biotechnology, DNA sequencing, DRT7, genome engineering, non-LTR-retroelements, retroviruses, RNA-dependent RNA polymerases, RNA sequencing, TGIRT-seq, UG10, Primase-Polymerase

## Abstract

Gene disruption analysis revealed that an *E. coli* PPRT protein, which has an N-terminal Primase-Polymerase (PrimPol) domain fused to a group II intron-like reverse transcriptase (RT) domain followed by a long C-terminal domain (CTD), contributes to a cellular oxidative DNA damage response in addition to its previously described function in phage defense. Biochemical analysis showed that the PrimPol domain has an error-prone DNA polymerase activity that enables read through of oxidation-induced DNA damage. Surprisingly, we found that the RT-like domain, in addition to synthesizing protein-primed DNAs for phage defense, has a 3’ to 5’ DNA exonuclease activity that functions in proofreading DNAs synthesized by the PrimPol domain. Extending these findings, we identified structural features that contribute to this proofreading activity, enabling us to associate it with both a group II intron-encoded and retroviral RT and suggesting general methods for incorporating proofreading activity into RTs.

## Introduction

Reverse transcriptases (RTs) are ancient enzymes that evolved from an RNA-dependent RNA polymerase (RdRP), likely during the transition from an RNA to DNA world^1–4^. Most impactful for evolution were prokaryotic mobile group II introns RTs, which promote both RNA splicing and group II intron mobility to unoccupied sites in DNA genomes^5–9^. Mobile group II introns and their RTs ultimately evolved into eukaryotic spliceosomal introns and key components of the eukaryotic splicing apparatus (U2, U4, and U5 snRNAs, and core spliceosomal protein Prp8) and in multiple steps into eukaryotic LINE-1 and other non-LTR-retrotransposon, telomerase, and retroviral RTs^10^. In parallel, bacterial mobile group II intron RTs diversified into numerous distinct and still largely unexplored subfamilies that evolved to perform different cellular functions, including phage defense by multiple mechanisms, RNA spacer acquisition in CRISPR-Cas systems, *de novo* synthesis of phage defense proteins, and DNA damage repair, a function also found recently for certain retroviral RTs^11–21^.

Mobile group II intron, retroplasmid, and human LINE-1 and other eukaryotic non-LTR-retrotransposon RTs, collectively termed non-LTR-retroelement RTs, remain structurally similar to each other as well as to RdRPs^4^. In addition to conserved RT sequence blocks RT1 to RT7 found in all RTs, non-LTR-retroelement RTs contain additional functionally important regions that are present in RdRPs but lost from retroviral RTs: an N-terminal extension (NTE) with an RT0 loop and extended regions (RT2a and RT3a) between conserved RT sequence blocks (Figure 1A)^1–4^. These additional regions have diversified in various ways that optimize bacterial RTs to perform different cellular functions^18,20,22,23^.

**Figure 1.**
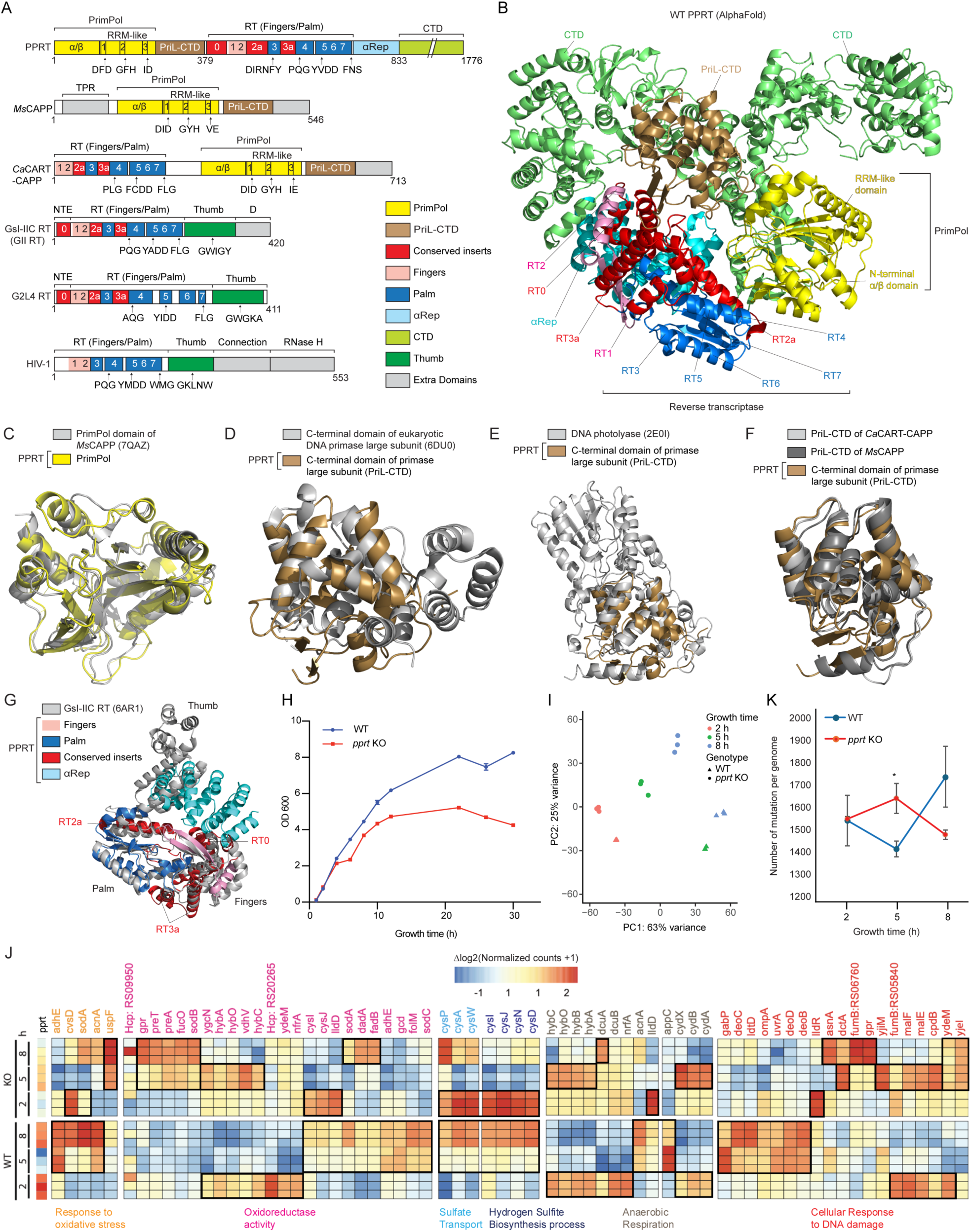
Characteristics and cellular function of PPRT. (A) Schematics comparing the *E. coli* SC366 PPRT protein with *Marinitoga* sp. 1137 *Ms*CAPP; *Caloramator australicus Ca*CART-CAPP; *Geobacillus stearothermophilus* group II intron GsI-IIC RT (denoted GII RT); a domesticated *Pseudomonas aeruginosa* group II intron-like RT (G2L4 RT) that functions in cellular DNA repair; and retrovirus HIV-1 RT. Protein regions are color-coded as indicated in the Figure. Abbreviations: CTD, C-terminal domain; NTE, N-terminal extension; TPR, Tetratricopeptide Repeat. (B) Three-dimensional model of PPRT predicted by AlphaFold 3. (C) AlphaFold 3 model of the PrimPol domain of PPRT (yellow) superimposed on *Marinitoga* sp. 1137 *Ms*CAPP (7QAZ) (gray). (D) AlphaFold 3 model of the PriL-CTD of PPRT (brown) superimposed on *Saccharomyces cerevisiae* DNA primase large subunit (6DU0) (gray). (E) AlphaFold 3 model of PriL-CTD of PPRT (brown) superimposed on *Sulfurisphaera tokodaii* str. 7 DNA photolyase (2E0I) (gray). (F) AlphaFold 3 model of PriL-CTD of PPRT (brown) superimposed on AlphaFold 3 model of PriL-CTD of *Caloramator australicus Ca*CART-CAPP and *Marinitoga* sp. 1137 *Ms*CAPP. (G) AlphaFold 3 model of the RT-like domain of PPRT with regions color-coded as in the panel A superimposed on *Geobacillus stearothermophilus* GsI-IIC RT (PDB: 6AR1; gray). (H) Growth curves of *E. coli* WT and *pprt* disruptant strains. Error bars represent standard deviation for three independent biological replicates. (I) Principal component analysis of gene expression differences identified by TGIRT-seq between the WT and *pprt* disruptant strains at different time points. Input gene read counts were normalized by a size factor determined by DESeq2. (J) Heatmaps comparing the expression of protein-coding genes in differentially expressed gene sets identified by geneset enrichment analysis in the *pprt* disruptant versus WT at different time points. Tiles are color coded by TGIRT-seq quantitation of the relative abundance of expressed RNAs compared to the row-wise mean value of log_2_ transformed DESeq2 normalized read counts. Black boxes in the heat map highlight the relative abundance of substantially over-expressed genes. (K) Mutation analysis. TGIRT-seq reads were used as input for VCF tools (vcftools.github.io) to quantitate mutations in the WT and *pprt* disruptant strains.

Numerous bacterial RTs identified by genome sequencing remain to be studied in detail. Here, we focused on an *Escherichia coli* PrimPol/RT fusion protein, which we denote PPRT. This protein has an N-terminal Primase-Polymerase (PrimPol) domain related to those of an archaea-eukaryotic primase superfamily that initiate DNA synthesis *de novo*, followed by a group II intron-like RT domain, and a long (~1,000 amino acid) C-terminal domain that is conserved in PPRTs but not found other proteins^17^. A previous study found that this PPRT protein, classified as Defense-associated RT type 7 (DRT7) and Unknown Group 10 (UG10), functions in phage defense, possibly via single-stranded (ss) DNA synthesis similar to DRT2, DRT9, and Class 1 UG/Abortive infection (Abi) RTs^12,17,24–29^. Surprisingly, we found that in addition to an analogous but not identical phage defense mechanism, the *E. coli* PPRT protein contributes to a cellular oxidative stress response by bypassing oxidation-induced 8-oxoguanine (8-oxo-dG) lesions and preferentially incorporating dC over dA at those sites. Even more surprisingly, we found that the PPRT protein has a 3’ to 5’ exonuclease activity that is associated with the RT-like domain and functions in proofreading DNAs synthesized by the PrimPol domain and that this proofreading activity could be transferred to a bacterial group II intron-encoded RT (*Geobacillus stearothermophilus* GsI-IIC RT) and a retroviral RT (HIV-1 RT) to increase their fidelity.

## Results

### Characteristics of the *pprt* gene and its encoded PPRT protein

The *E. coli* SC366 PPRT protein, the focus of this study, is encoded within a bacterial defense island surrounded by genes encoding proteins that may mobilize this DNA segment, including a transposase, genes associated with conjugative plasmids (*trb*L, *trb*J, and *tra*J), and a prophage (CP4-57; Figure S1A). A BLASTP search identified several hundred genes encoding proteins that have strong homology to this PPRT protein, with most found in γ-proteobacteria and enteric bacteria, including *E. coli*, *Salmonella* spp., *Klebsiella* spp., *Vibrio* spp., and *Pseudomonas* spp. (Figure S1B). These PPRTs are typically large proteins (1,700 to 1,900 amino acids (aa)), consisting of an N-terminal PrimPol domain (~400 aa) followed by an RT-like domain (~300 aa), and a long C-terminal domain (~1,000 aa), which lacks sequence homology to any other known protein but is conserved in PPRTs (Figure 1A).

AlphaFold 3.0 models of full-length *E. coli* PPRT (Figure 1B) and its individual domains were used as structural queries for Foldseek searches against the Protein Data Bank (PDB)^30,31^. The full-length PPRT model showed high local confidence, with most residues showing predicted Local Distance Difference Test (pLDDT) scores >70 with many >90, and a predicted Template Modeling (pTM) score of 0.79. Protein Data Bank (PDB) structures corresponding to the top Foldseek hits were structurally aligned in PyMOL using the cealign command, which involves combinatorial extension (CE) algorithm^32,33^. Root Mean Square Deviation (RMSD) values were calculated over alpha-carbon (Cα) pairs matched by the CE algorithm.

The top-ranked match for the N-terminal PrimPol domain of PPRT was *Ms*CAPP (PDB ID: 7QAZ)^34^, a CRISPR-associated PrimPol from *Marinitoga* sp. 1137 (Figure 1C; RMSD 2.4 Å). The PPRT PrimPol domain includes an initial α/β segment and an RNA Recognition Motif-like (RRM-like) segment with three conserved active site motifs: motif I (DFD), motif II (GFH), and motif III (ID) (Figure 1A). Motifs I and III contain aspartate residues that bind catalytic metal ions, and motif II has a nucleotide binding motif^35,36^. The PrimPol domain is followed by a primase large subunit C-terminal domain (PriL-CTD), with the top-ranked matches *Saccharomyces cerevisiae* PriL-CTD (Figure 1D; RMSD 4.6 Å) and *Sulfurisphaera tokodaii* str. 7 DNA photolyase (Figure 1E; RMSD 4.8 Å). Eukaryotic PriL-CTD binds a single-stranded (ss) DNA template and helps position initial ribonucleotides for RNA primer synthesis, while the DNA photolyase binds damaged DNA and positions pyrimidine dimers for repair^37^. PriL-CTD of PPRT is also similar to PriL-CTD found in AlphaFold3 predicted models of *Ms*CAPP (Figure 1F; RMSD 3.4 Å) and *Caloramator australicus Ca*CART-CAPP (Figure 1F; RMSD 3.1 Å), a CRISPR-associated RT/PrimPol fusion protein in which the RT domain precedes the PrimPol domain.

The PPRT RT-like domain fused to PriL-CTD corresponds to the fingers and palm regions of group II intron-encoded RTs with a predicted structure similar to that of GsI-IIC RT (Figure 1G; RMSD 4.2 Å). It contains RT1-7 regions present in all RTs plus an NTE with an RT0 loop and RT2a and RT3a between the conserved RT sequence blocks present in group II intron-encoded and other non-LTR-retroelement RTs (Figure 1A)^4^. Its conserved active site motif in RT5 is YVDD with D626 and D627 corresponding to conserved aspartate residues that bind catalytic divalent metal ions, and D526 in RT3 corresponding to a third conserved aspartate that contributes to binding divalent cations in other RTs^4^. The PPRT RT6 and 7 motifs differ from those found in other RTs as described further below.

Downstream of the RT-like domain, the PPRT protein has a predicted α-helical repeat (αRep) region analogous to those found in AbiA, AbiK, and Abi-P2 RTs, which function in stabilizing ssDNA products that are required for the phage defense capabilities of Class 1 UG/Abi RTs^17,24,25^. This αRep region replaces the canonical RT thumb domain (Figure 1G), creating a confined space between the αRep and fingers subdomains similar to the architecture observed in AbiA, AbiK, and Abi-P2 RTs^25^. The αRep domain is followed by a long, conserved C-terminal domain (CTD; 944 amino acids) predicted by AlphaFold 3.0 to be comprised largely of sets of α-helices that fold back over the extended PrimPol and RT domains (Figure 1B).

### The PPRT protein functions in oxidative damage response *in vivo*

To explore the biological function of PPRT, we disrupted the wild-type (WT) *pprt* gene in *E. coli* SC366 by inserting a programmable RNA-guided mobile group II intron (targetron)^38,39^ into a region of the gene encoding the RT domain (nucleotide 1218s; Figure S1A). After confirming targetron insertion at the desired site by PCR and Southern hybridization (Figures S1C and S1D), we cured the targetron expression plasmid to impede RNA splicing of the inserted targetron and confirmed a single targetron insertion at the desired site by whole-genome sequencing. The growth curves of the WT and *pprt* disruptant strains in liquid culture were similar for up to 2 h, after which growth of the disruptant slowed for up to 5 h, and then resumed at a fast rate but with an earlier entry into stationary phase than the wild-type strain (Figure 1H).

Based on the growth curves, we comprehensively profiled protein-coding and non-coding RNAs in the WT and the *pprt* disruptant strains at 2, 5, and 8 h by using a Thermostable Group II Intron Reverse Transcriptase sequencing (TGIRT-seq) protocol that enables parallel quantitative analysis of chemically fragmented mRNAs and long non-coding (lnc) RNAs (>200 nt) together with intact small non-coding (snc) RNAs (≤200 nt)^40^. Principal component analysis of triplicate TGIRT-seq datasets for the WT and disruptant strains confirmed reproducible gene expression differences between these strains at each time point attributable to changes in *pprt* gene expression and growth rate (PC2 and PC1, respectively; Figure 1I). Volcano plots comparing the relative abundance of transcripts between WT and the disruptant identified ~60 genes at 2 h increasing to >200 genes at 5 h and 8 h that were strongly differentially expressed (log_2_ Fold change >3 or <-3, adjusted p-values <0.00001; Figures S1E-S1G). The expression levels of most sncRNAs were similar between the WT and disruptant strains, with the exception of several tRNAs and RNase P, which were significantly over-expressed in the disruptant at 8 h, and antisense sRNA RprA, an environmental stress response regulator^41^, which was substantially under-expressed in the disruptant at later time points (Figures S1F and S1G).

To analyze differences in the expression of functionally related protein-coding genes at different time points, we performed gene set enrichment analysis using ShinyGO^42^, a method that enables analysis of differences due to loss of function of the *pprt* gene, growth time, and the interaction between these two parameters (Figure S1H and Methods). This analysis found significant often reciprocal differences between the WT and the disruptant at different time points in the gene sets Response to Oxidative Stress, Oxidoreductase Activity, Sulfate Transport, Hydrogen Sulfite Biosynthesis Process, Anaerobic Respiration, and Cellular Response to DNA Damage (Figure 1J).

In the WT strain, the *pprt* gene was expressed at the highest level at 2 h, lower levels at 5 h, and moderately higher levels at 8 h. In addition to *pprt*, other genes expressed at high levels in WT cells at 2 h included subsets that contribute to oxidoreductase activity, anaerobic respiration, and cellular response to DNA damage (Figure 1J). At 5 and 8 h, lower expression levels of the *pprt* gene in the WT strain were correlated with higher levels of other genes that were suggestive of increases in response to oxidative stress, oxidoreductase activity, and cellular response to oxidative DNA damage (Figure 1J). By contrast, expression of the above gene sets in the *pprt* disruptant showed differences suggesting a higher level of response to oxidative stress at 2 h, more sustained expression levels of anaerobic respiration genes at 5 and 8 h, and a more sustained induction of different sets of cellular response to DNA damage genes at 5 and 8 h (Figure 1J).

Collectively, these findings suggested that PPRT contributes to a cellular response to oxidative stress and DNA damage, possibly resulting from growth in shaking liquid culture flasks (see Methods). This response was more effective in the WT than the disruptant at 2 h, leading to a decrease in PPRT expression at 5 h, followed by an increase at 8 h. This scenario was supported by a significantly lower frequency of genomic mutations identified by analysis of the TGIRT-seq datasets in the WT strain at 5 h followed by an increased frequency at 8 h and a higher frequency of mutations in the *pprt* disruptant at 5 h followed by a decrease at 8 h when the disruptant began an early entry into stationary phase (Figure 1K and Table S1).

### PPRT has both DNA-dependent DNA polymerase and DNA exonuclease activities

To investigate the biochemical activities of the WT PPRT protein, we began by carrying out primer extension assays with purified protein and 3’-inverted dT (3’-Inv(dT))-blocked 50-nt DNA or RNA templates of the same sequence annealed to a 5’-^32^P-labeled 20-nt DNA primer. Reactions were initiated by adding 1 mM dNTPs (an equimolar mixture of 1 mM dATP, dCTP, dGTP, and dTTP) in reaction medium containing different concentrations of Mg^2+^ and/or Mn^2+^ (Figures 2A-2C) and incubated at 37°C for 30 min. These assays showed that the primer extension activity of WT PPRT with a DNA template was favored at higher Mg^2+^ concentrations and more so at higher Mn^2+^ concentrations (Figures 2A and 2C), but was at most marginal with an RNA template under all conditions tested (Figure 2B).

**Figure 2.**
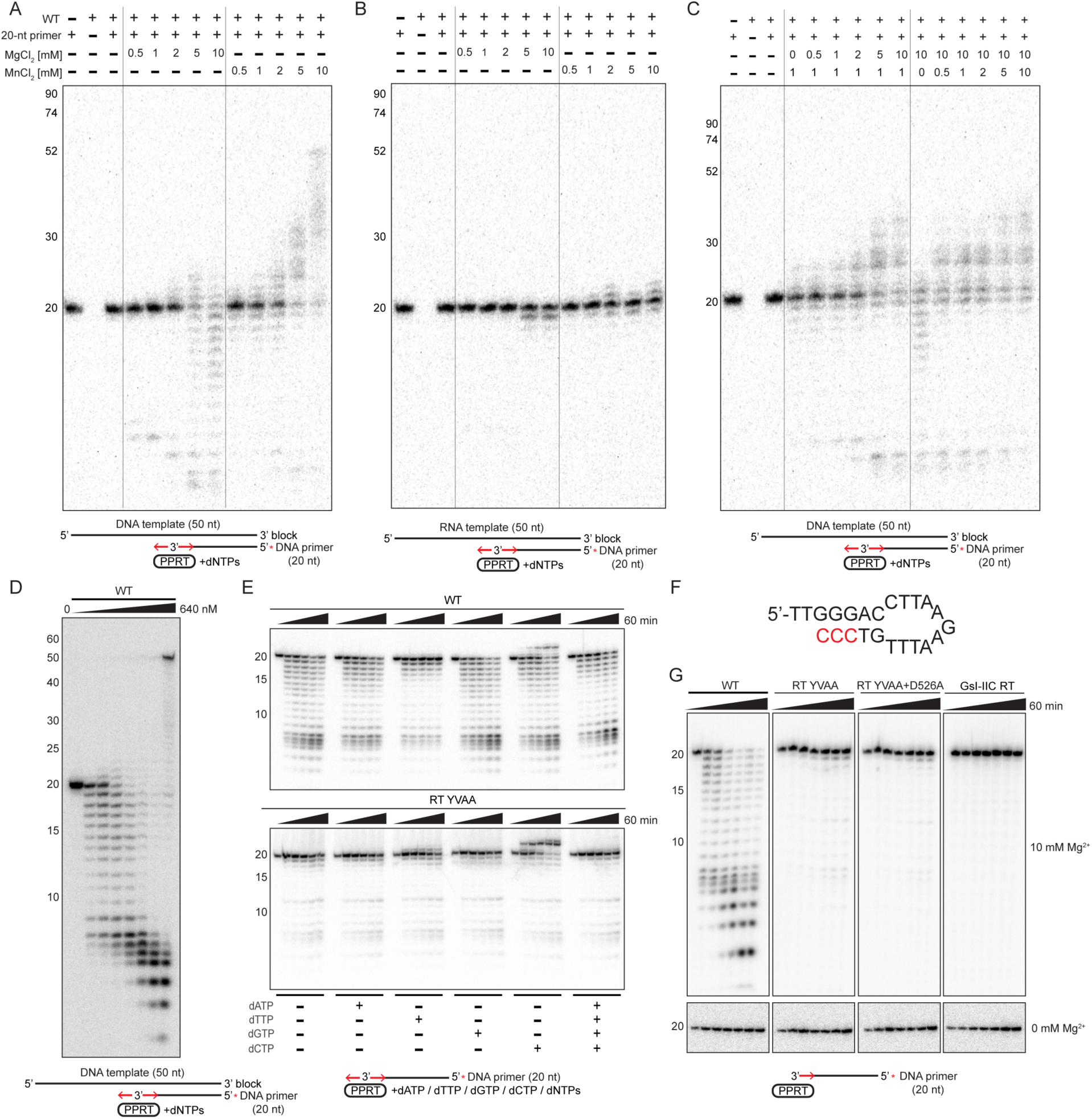
PPRT has both DNA-dependent DNA polymerase and DNA exonuclease activities. (A and B) Primer extension of WT PPRT in reaction medium containing different concentrations of MgCl_2_ or MnCl_2_ assayed with a 3’-inverted dT (3’-Inv(dT))-blocked 50-nt DNA (A) or RNA (B) template annealed to a 5’-^32^P-labeled (red asterisk) 20-nt DNA primer. Assays were done as described in Methods by incubating 100 nM WT PPRT with 20 nM of template-primer substrate in a reaction medium containing 10 mM NaCl, different concentrations of MgCl_2_ and MnCl_2_, 20 mM Tris-HCl pH 7.5, and 1 mM dNTPs (an equimolar mix of 1 mM dATP, dCTP, dGTP, and dTTP) at 37°C for 30 min. DNA products were analyzed in a denaturing 17% polyacrylamide gel against 5’-labeled oligonucleotide size markers in a parallel lane. (C) Divalent cation dependence of primer extension activity of WT PPRT with different combinations of MgCl_2_ and MnCl_2_ assayed as in panels A and B. (D) Protein-concentration dependence of WT PPRT primer extension activity. Reactions were done as in panel A in reaction medium containing 10 mM MgCl_2_, and 1 mM MnCl_2_ for 30 min with different concentrations of PPRT (0, 10, 20, 40, 80, 160, 320, 640 nM). (E) Competing snapback DNA synthesis and exonuclease assays of WT and YVAA mutant PPRT proteins. Reactions were done as in panel D with 100 nM PPRT and 20 nM 5’-^32^P-labeled (red asterisk) 20-nt DNA oligonucleotide substrate in reaction medium containing 10 mM MgCl_2_, 1 mM MnCl_2_, and 1 mM of each dNTP indicated below the autoradiogram for times up to 60 min. (F) Predicted snapback DNA structure for the extended terminal transferase product seen with dCTP in panel E with three added 3’ dC residues (red) complementary to three 5’-proximal dG residues. (G) Exonuclease assays of WT and mutant YVAA, YVAA + D526A PPRTs and GsI-IIC RT. Reactions were done as in panel E with a 5’-^32^P-labeled 20-nt DNA primer for times up to 60 min. The bottom panel shows exonuclease assays of the same proteins in the absence of Mg^2+^ in reaction medium containing 100 nM enzyme and 25 mM EDTA at 37°C for times up to 60 min.

Surprisingly, these assays also showed that PPRT has a Mg^2+^-favored 3’ to 5’ DNA exonuclease activity with addition of Mn^2+^ increasing primer extension activity and decreasing exonuclease activity, suggesting competition between these activities (Figures 2A and 2C). Assays with increasing concentrations of WT PPRT (0 to 640 nM) in reaction medium containing 10 mM Mg^2+^ and 1 mM Mn^2+^ for 1 h showed progressive increases in both primer extension and exonuclease activity, with exonuclease activity dominant at the lowest PPRT concentration and a full-length 50-nt DNA synthesized at the highest PPRT concentration (Figure 2D), also indicative of competition of these activities with primer extension activity favored by higher PPRT concentrations.

Additional assays of WT PPRT with only a 5’-labeled 20-nt DNA primer in reaction medium containing 10 mM Mg^2+^ and 1 mM Mn^2+^ showed a predominant 3’ to 5’ exonuclease activity in the presence of each individual or all four dNTPs (Figure 2E, top panel). The reaction with 1 mM dCTP showed an additional higher molecular weight band that likely reflects snapback DNA synthesis that adds three 3’-dC residues complementary to three dG residues near the 5’ end of the 20-nt oligonucleotide template (Figure 2E and 2F). A PPRT mutant with YVAA instead of YVDD at the RT active site produced higher levels of a similar snapback product with dCTP, indicating that it was synthesized by the PrimPol domain (Figures 2E and 2F). This snapback product was detected in the presence of dCTP, but not any of the other or all four dNTPs, the latter likely reflecting competition between snapback DNA synthesis and protein-primed phage defense DNA synthesis that preferentially consumes dTTP and dATP (see below). Notably, the YVAA mutant had low residual exonuclease activity, which was decreased further in the YVAA + D526A mutant, which lacked all three conserved RT active site aspartate residues, indicating that this is an additional activity of the RT-like domain (Figure 2G, top panel). The low residual exonuclease activity was not detected in reaction medium lacking Mg^2+^ ions (Figure 2G, bottom panel), suggesting that it reflected residual Mg^2+^ binding at a proximate or interacting site, supported by a finding below. Group II intron-encoded GsI-IIC RT, which has YADD at its active site, showed no comparable exonuclease activity (Figure 2G). Collectively, these findings suggested that the PPRT protein’s PrimPol domain has a Mn^2+^-favored primer extension activity and that its RT-like domain has a Mg^2+^-favored 3’ to 5’ DNA exonuclease activity.

### Mutational analysis of primer extension and DNA exonuclease activities and the contribution of the C-terminal domain to these activities

For further analysis, we compared the primer extension and exonuclease activities of WT and mutant PPRT proteins with point mutations in PrimPol or RT-like domain active-site residues (Figure 3A). These reactions were done as time courses with the same 3’-blocked 50-nt DNA template and an annealed 5’-labeled 20-nt DNA primer in a reaction medium containing 10 mM MgCl_2_ and 1 mM MnCl_2_ (Figures 3B and 3C). Under these conditions, WT PPRT primer extension activity competed with its 3’ to 5’ exonuclease activity, indicated by a time-dependent increase in longer, radiolabeled products in the presence but not in the absence of dNTPs (Figures 3B and 3C). The PrimPol domain D68A, D70A, and D68A+D70A mutants, which completely lacked primer extension activity, retained 3’ to 5’ exonuclease activity, indicated by the progressive shortening of the labeled annealed primer in the presence or absence of dNTPs (Figures 3B and 3C, respectively). By contrast, the RT-like domain YVAA mutant, which retained primer extension activity, showed greatly decreased exonuclease activity, enabling it to produce a full-length 50-nt DNA product in the presence of dNTPs (top band in Figure 3B). The mutant D68A+D70A+YVAA, in which catalytic aspartate residues in both the PrimPol and RT-like domain active sites were replaced by alanines, showed minimal primer extension activity and lower DNA exonuclease activity than the YVAA mutant, suggesting the PrimPol active site as possible source of residual Mg^2+^ binding for DNA exonuclease activity in the YVAA mutant (Figures 3B and 3C).

**Figure 3.**
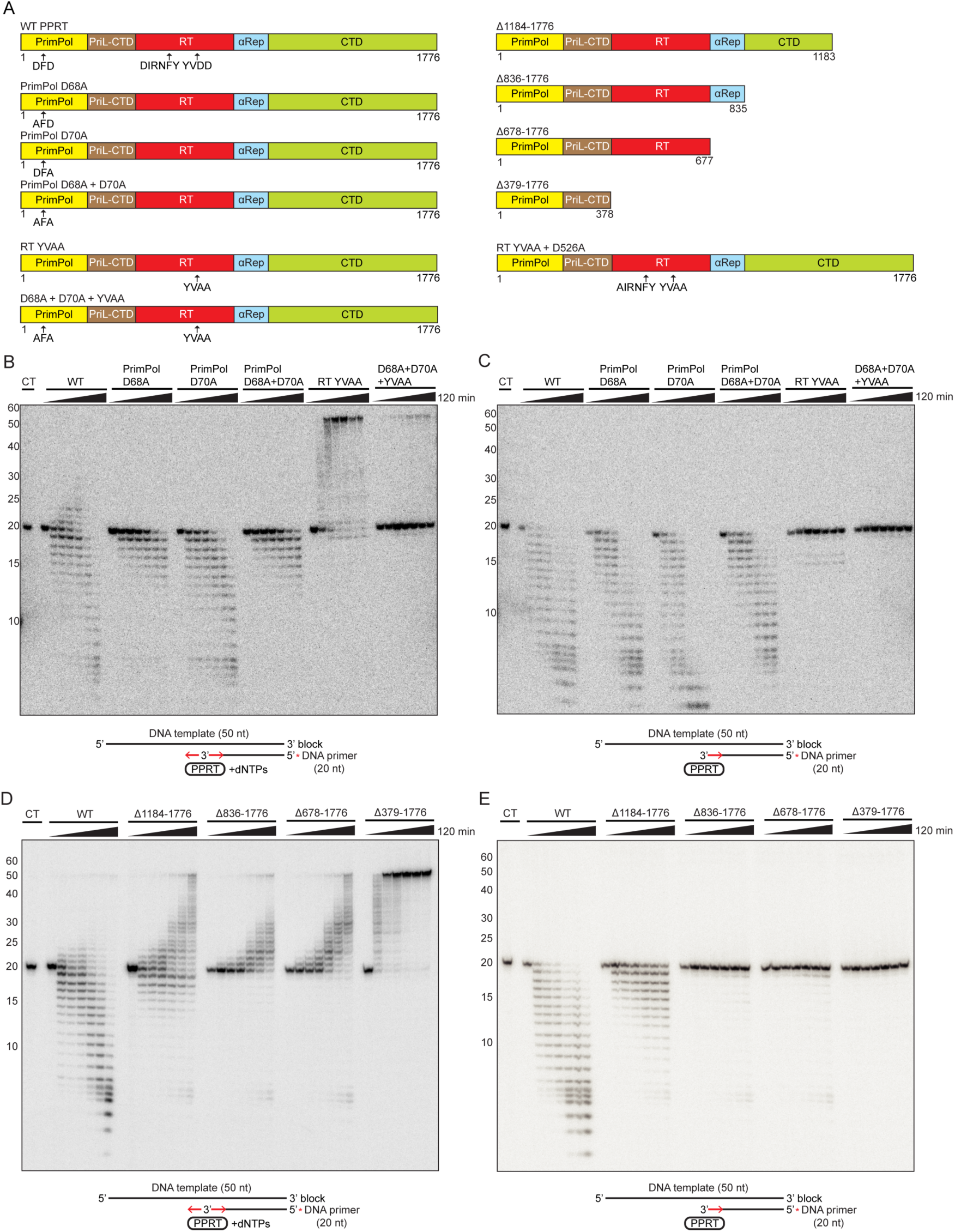
The PPRT PrimPol domain has primer extension activity while the RT domain has 3’ to 5’ DNA exonuclease activity. (A) Schematics of WT and mutant PPRTs. (B) Primer extension assays of WT and mutant PPRTs using a 3’-Inv(dT)-blocked 50-nt DNA template and a 5’-^32^P-labeled (red asterisk) 20-nt DNA primer. Reactions were done at 37°C for times up to 120 min. CT (control) was tested without enzyme under identical conditions, with an incubation time of 120 min. (C) Exonuclease assay of WT and mutant PPRTs with the same 3’-Inv(dT)-blocked 50-nt DNA template and a 5’-^32^P-labeled (red asterisk) 20-nt DNA primer used in panel B. Reactions were done as in panel B in the absence of added dNTPs at 37°C for times up to 120 min. CT (control) was tested without enzyme under identical conditions, with an incubation time of 120 min. (D) Primer extension assays of WT and C-terminal deleted PPRTs done as in panel B. (E) Exonuclease assays of WT PPRT and C-terminal deletion mutants done as in panel C. All of the above reactions were done in reaction medium containing 100 nM enzyme, 10 mM NaCl, 10 mM MgCl_2_, 1 mM MnCl_2_, and 20 mM Tris-HCl pH 7.5, with the DNA products analyzed in a denaturing 17% polyacrylamide gel against 5’-labeled oligonucleotide size markers in a parallel lane (see Methods).

To investigate the function of the long C-terminal domain of PPRT, we performed similar primer extension and exonuclease assays comparing WT PPRT with mutant proteins having progressively long C-terminal deletions: Δ1184-1776, Δ836-1776, Δ678-1776, and Δ379-1776, the latter corresponding solely to the PrimPol domain (Figures 3A, 3D, and 3E). Interestingly, the mutants with longer C-terminal deletions had progressively higher primer extension activity, generating more full-length 50-nt cDNA products, indicating lower competing DNA exonuclease activity (Figure 3D). This was confirmed by assays in the absence of dNTPs, where the shortest C-terminal truncation mutant (Δ1184-1776) retained moderate exonuclease activity, while proteins with longer C-terminal truncations (Δ678-1776 and Δ836-1776), which retained the complete RT-like domain, nevertheless had very low exonuclease activity (Figure 3E). Finally, the Δ379-1776, which retained only the PrimPol domain, had by far the highest primer extension activity and completely lacked any residual exonuclease activity (Figure 3D). Collectively, these findings indicated (i) that the PrimPol domain has primer extension activity and the RT-like domain has a competing 3’ to 5’ exonuclease activity that can access the 3’ end of an annealed primer, and (ii) that the PPRT C-terminal domain contributes to exonuclease activity, possibly by helping to establish correctly structured or enclosed regions of the RT-like domain.

### The RT-like domain of PPRT also has a protein-primed DNA synthesis activity

PPRT has been classified as a DRT7/UG10 protein related to Class 1 UG/Abi and Class 2 UG/Abi DRT2 and DRT9 RTs that carry out ssDNA synthesis for phage defense^17,25,26,29^. Similar to Class 1 UG/Abi RTs, PPRT has a C-terminal α-helical domain in place of a conventional RT thumb domain (Figure 4A)^17^. While DRT2 and DRT9 use associated non-coding (nc) RNAs as templates for ssDNA synthesis^26–29^, PPRT lacks predicted neighboring ncRNAs (Figure S1A)^17^. Although we found that the PPRT RT domain has exonuclease activity, given that some Class 1 UG/Abi RTs (*e.g.*, AbiA, AbiK, and Abi-P2) achieve phage resistance via template-independent ssDNA synthesis and that PPRT has been shown to provide phage resistance^17,24,25^, we tested whether PPRT has a similar ssDNA synthesis activity.

**Figure 4.**
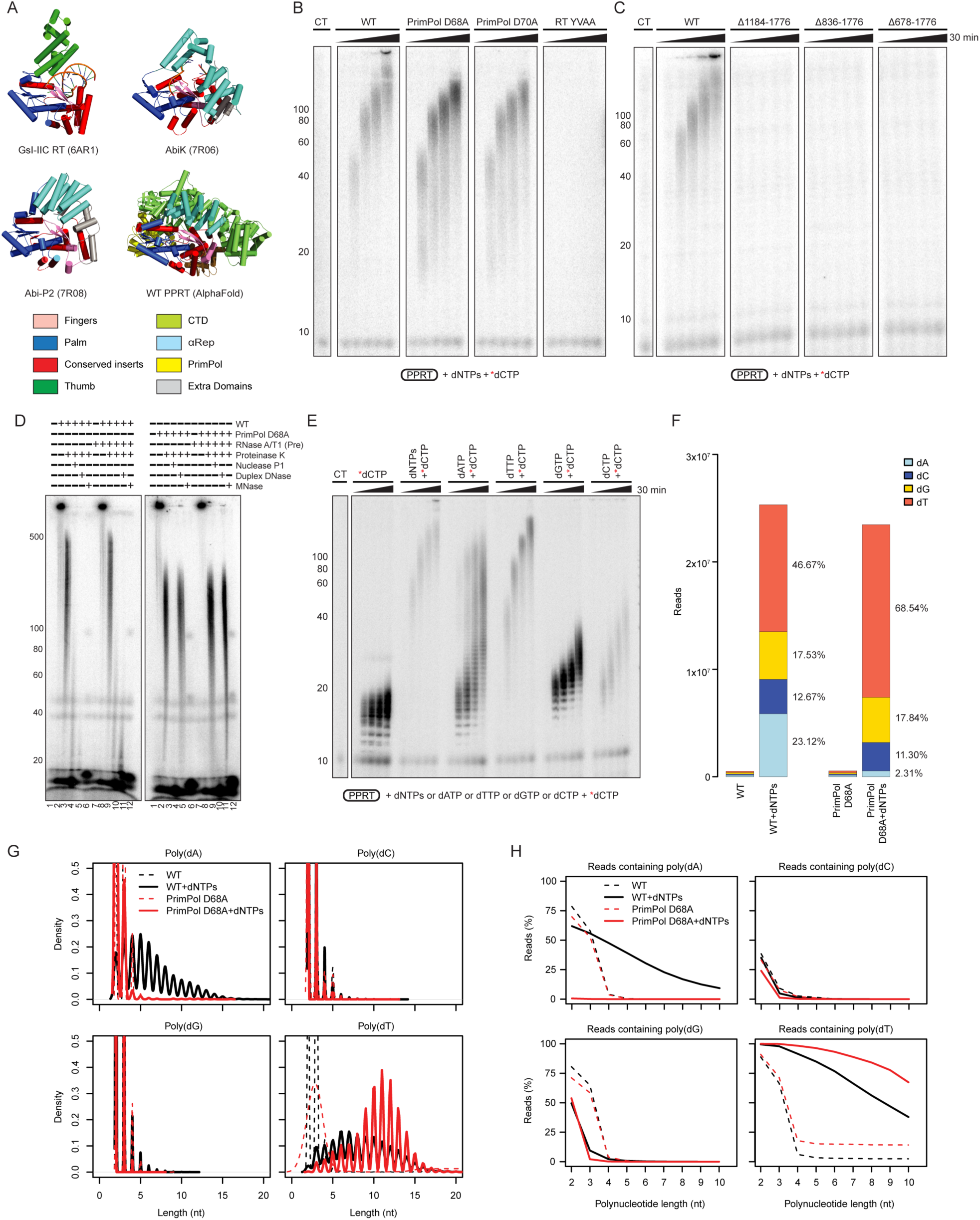
PPRT carries out protein-primed DNA synthesis. (A) Structural comparison of GsI-IIC RT (PDB: 6AR1), AbiK (PDB: 7R06), Abi-P2 (PDB: 7R08), and WT PPRT (AlphaFold3). (B) Template-independent DNA polymerization of WT and mutant PPRTs using 50 μM dNTPs plus 83 nM [α-^32^P]-dCTP (red asterisk) in the absence of DNA templates and primers in reaction medium containing 200 nM enzyme, 10 mM NaCl, 10 mM MgCl_2_, 1 mM MnCl_2_, and 20 mM Tris-HCl pH 7.5 at 37°C for times up to 30 min. The DNA products were analyzed in a denaturing 17% polyacrylamide gel against 5’-labeled oligonucleotide size markers in a parallel lane (see Methods). CT (control) was tested without enzyme under identical conditions, with an incubation time of 30 min. (C) Template-independent DNA polymerization by WT and PPRT C-terminal deletion mutants done as in panel B. (D) Template-independent DNA polymerization reactions in which WT or PrimPol domain D68A mutant PPRT proteins (200 nM) were pre-incubated with or without RNase A/T1 prior to addition of 50 μM dNTPs plus 83 nM [α-^32^P]-dCTP (red asterisk) at 37°C for 30 min. After treatment with thermolabile Proteinase K and heat inactivation at 55°C for 30 min, samples were digested with nuclease P1, Duplex DNase, or micrococcal nuclease at 37°C for 20 min (see Methods). The DNA products were analyzed in a denaturing 6% polyacrylamide gel against 5’-labeled oligonucleotide size markers in a parallel lane (see Methods). (E) Template-independent DNA polymerization of WT PPRT done as in panel B using 50 μM of all four dNTPs or a single dNTP plus 83 nM [α-^32^P]-dCTP (red asterisk). CT (control) was tested without enzyme with 83 nM [α-^32^P]-dCTP under identical conditions, with an incubation time of 30 min. (F) Composition of DNAs detected in WT and PrimPol D68A (200 nM) incubated for 30 min without dNTPs (WT, PrimPol D68A, respectively) or with 50 μM dNTPs (WT+dNTPs, PrimPol D68A+dNTPs, respectively) in the absence of [α-^32^P]-dCTP. The stacked bar graphs show percentages of each nucleotide (dA, dC, dG, and dT) measured by Illumina MiSeq sequencing (see Methods). (G) Density plots showing the length distribution of poly(dN) runs ≥2 nt (N = A, C, G, or T) in DNAs synthesized by reactions done as in panel F. For WT + dNTPs, 39% of the reads contained dA runs ≥5 nt, while for PrimPol D68A + dNTPs, only 0.4% of the reads contained dA runs ≥5 nt (top left panel). Both enzymes exhibited minimal incorporation of dCTP or dGTP, with <0.5% of reads containing dC or dG runs ≥5 nt (top right and bottom left panels. For WT PPRT, 84% of reads contained poly(dT) runs ≥5 nt, while for the PrimPol D68A mutant, 96% of reads contained poly(dT) runs ≥5 nt (bottom right panel). (H) Plots showing the polynucleotide length distribution of poly(dA), poly(dC), poly(dG), and poly(dT) runs in DNAs synthesized by reactions done as in panel F.

In an initial experiment, we incubated WT and mutant PPRTs with 50 μM dNTPs and trace (83 nM) [α-^32^P]-dCTP in reaction medium containing 10 mM Mg^2+^ and 1 mM Mn^2+^ and found that WT PPRT efficiently synthesized ^32^P-labeled DNAs without an added primer or template (Figure 4B). WT PPRT generated 30-60 nt products within 1 min, with products extending well beyond 100 nt after 30 min. While the PrimPol D68A and D70A mutants retained this DNA synthesis activity, the RT-like domain active site-mutant YVAA showed no such activity (Figure 4B), indicating that the RT active site catalyzes this additional activity that may take priority over its exonuclease activity when triggered for phage defense.

Building on our previous findings that C-terminal domain deletion mutants significantly impacted RT-like domain exonuclease activity (Figures 3D and 3E), we did additional template-independent DNA synthesis assays comparing WT PPRT with C-terminal deletion mutants Δ1184-1776, Δ836-1776, and Δ678-1776, the latter completely lacking the αRep and CTD and containing only the PrimPol and RT-like domain. All three mutants completely lacked template-independent DNA synthesis activity (Figure 4C), including the mutant protein with the shortest C-terminal truncation (Δ1184-1776), which retained moderate 3’ to 5’ exonuclease activity (Figure 3E). These findings indicated that the C-terminal domain contributed to non-templated DNA synthesis, most likely by helping to establish correctly structured or enclosed regions of the RT-like domain.

Abi (*e.g.*, AbiA, AbiK, and Abi-P2) and DRT9 RTs initiate ssDNA synthesis for phage resistance by a protein-priming mechanism using an amino acid residue containing an OH group to which the newly synthesized ssDNA can remain covalently attached^24–26,29^. DRT9 produces poly(dA)-rich ssDNAs using tyrosine residues in its C-terminal tail as protein-priming sites^26,29^. Indicative of protein priming, similar experiments confirmed that WT PPRT likewise synthesizes ^32^P-labeled DNAs that remain covalently attached to the PPRT protein and whose synthesis is inhibited by the YVAA mutation at the RT-like domain active site (Figure S2A).

To test if PPRT synthesizes ssDNAs similarly to those generated by DRT2 and DRT9 by reiterative copying of an RNA template, we preincubated WT and D68A mutant PPRTs with or without RNase A/T1, carried out DNA synthesis assays as above with 50 μM dNTPs and trace (83 nM) [α-^32^P]-dCTP, and then incubated the products with or without thermolabile proteinase K, nuclease P1, Duplex DNase, or micrococcal nuclease (MNase). As shown in Figure 4D, RNase pre-treatment had no effect on DNA synthesis, consistent with lack of an RNA template, and unless treated with Proteinase K, most of the ^32^P-labeled products remained in the well at the top of the gel, suggesting attachment to the PPRT protein via protein priming.

Notably, the labeled DNA products released by Proteinase K treatment ranged up to 500 nt for WT PPRT (Figure 4D, left panel), while those synthesized by PrimPol D68A mutant were shorter (Figure 4D, right panel), suggesting that, although the RT domain was required to initiate synthesis of these DNAs, the PrimPol domain contributed to the synthesis of longer DNAs. Treatment of the DNA products released after proteinase K digestion with nuclease P1 (which degrades ssDNA), Duplex DNase (which degrades double strand (ds) DNA), and MNase (which degrades both ssDNA and dsDNA) resulted in leftover products ranging from 20 to 80 nt for nuclease P1, 20 to 30 nt for Duplex DNase, and no detectable products for MNase (Figure 4D, left panel lanes 4, 5, 6, respectively), suggesting that WT PPRT generates both single-stranded and double-stranded DNAs. By contrast, DNA products synthesized by PrimPol mutant D68A were completely digested with nuclease P1 or MNase and were resistant to the Duplex DNase (Figure 4E, right panel, lanes 10-12), indicating that they were now limited to ssDNAs. Collectively, these findings indicated that the RT-like domain initiates synthesis of ssDNAs by protein priming, as does Abi and other DRTs^26,29^, and that the PrimPol domain functions in generating dsDNA products by copying regions of the initial ssDNA synthesized by the RT-like domain.

Further DNA synthesis assays with a single dNTP (dATP, dTTP, dGTP, or dCTP) and trace amounts of [α-^32^P]-dCTP showed different extension patterns with each dNTP, suggesting largely random, non-templated incorporation with the longest labeled products obtained with dTTP followed by dATP, dCTP, and dGTP (Figure 4E). Illumina MiSeq sequencing of DNAs synthesized by WT PPRT with an equimolar mixture of all four dNTPs confirmed that products synthesized by the WT enzyme had random sequences with preferences for dT>dA>dG>dC residues, while DNAs synthesized by PrimPol D68A had higher proportions of dT residues, similar levels of dC and dG residues, and very low levels of dA residues (Figure 4F). These findings indicated that the RT-like domain preferentially synthesized ssDNAs with dT residues, while the PrimPol domain preferentially contributed dA residues by copying the dT-rich ssDNA, resulting in partially dsDNAs. Sequencing confirmed that DNAs synthesized by WT PPRT had discrete runs of dT and dA residues, while those synthesized by the PrimPol D68A mutant had a higher proportion of dT runs and lacked dA runs (Figures 4G, S2B, and S2C). Both the WT and PrimPol D68A mutant enzymes exhibited minimal incorporation of dC or dG residues (Figure 4F). For WT PPRT, runs of dT residues were enriched at the 5’ terminus and in the middle portion of the reads, while runs of dA residues were enriched only in the middle portion (Figures S2D and S2E). Collectively, these findings indicated that the RT domain initiates protein-primed DNA synthesis of ssDNAs that contain runs of dT residues, and the PrimPol domain copies these DNAs to add double-stranded regions with runs of dA residues. The latter could be done either by *de novo* initiation, a known activity of PrimPol proteins^35,43^, or by snapback DNA synthesis of dissociated ssDNAs with a 3’ OH, the latter an activity of the PrimPol domain found in Figure 2E. The synthesis of dsDNAs could potentially contribute to phage defense by amplifying dNTP depletion or possibly by a mechanism different than ssDNAs.

### PPRT exonuclease activity increases fidelity

The AlphaFold model of PPRT bound to Mg^2+^ ions indicated that the active sites of the PrimPol and RT-like domains are separated by ~46 Å (Figure 5A), within the range observed in canonical DNA polymerases that have a functionally coordinated proofreading exonuclease activity at a separate active site^44,45^. To investigate if PPRT’s exonuclease activity functions in proofreading DNAs synthesized by the PrimPol domain, we performed fidelity assays comparing primer extension reactions using a 5’-labeled 20-nt primer and 50-nt 3’-blocked template strands that are either fully complementary to the primer or have one or two mismatches at its 3’ end and adding either dTTP or dATP for initiation of DNA synthesis (Figure 5B). For reactions with dTTP the tested enzyme is expected to copy the two upstream dA residues in the template resulting in a 22-nt DNA product and then stop, with any extension beyond that point reflecting misincorporation at non-complementary DNA template residues. For reactions with dATP, any extension of the primer would indicate misincorporation. The reactions were done with increasing concentrations of dTTP or dATP (1 μM to 1 mM) to test competition between primer extension and DNA exonuclease activity. This experimental design allowed us to test (i) whether RTs could utilize a primer with a mismatched 3’ nucleotide, a capability typically absent in polymerases that lack a proofreading exonuclease activity that would remove the primer’s mismatched 3’ nucleotide^46,47^, and (ii) whether the exonuclease activity enhances fidelity by minimizing incorporation of non-complementary nucleotides during primer extension^43,45,48^.

**Figure 5.**
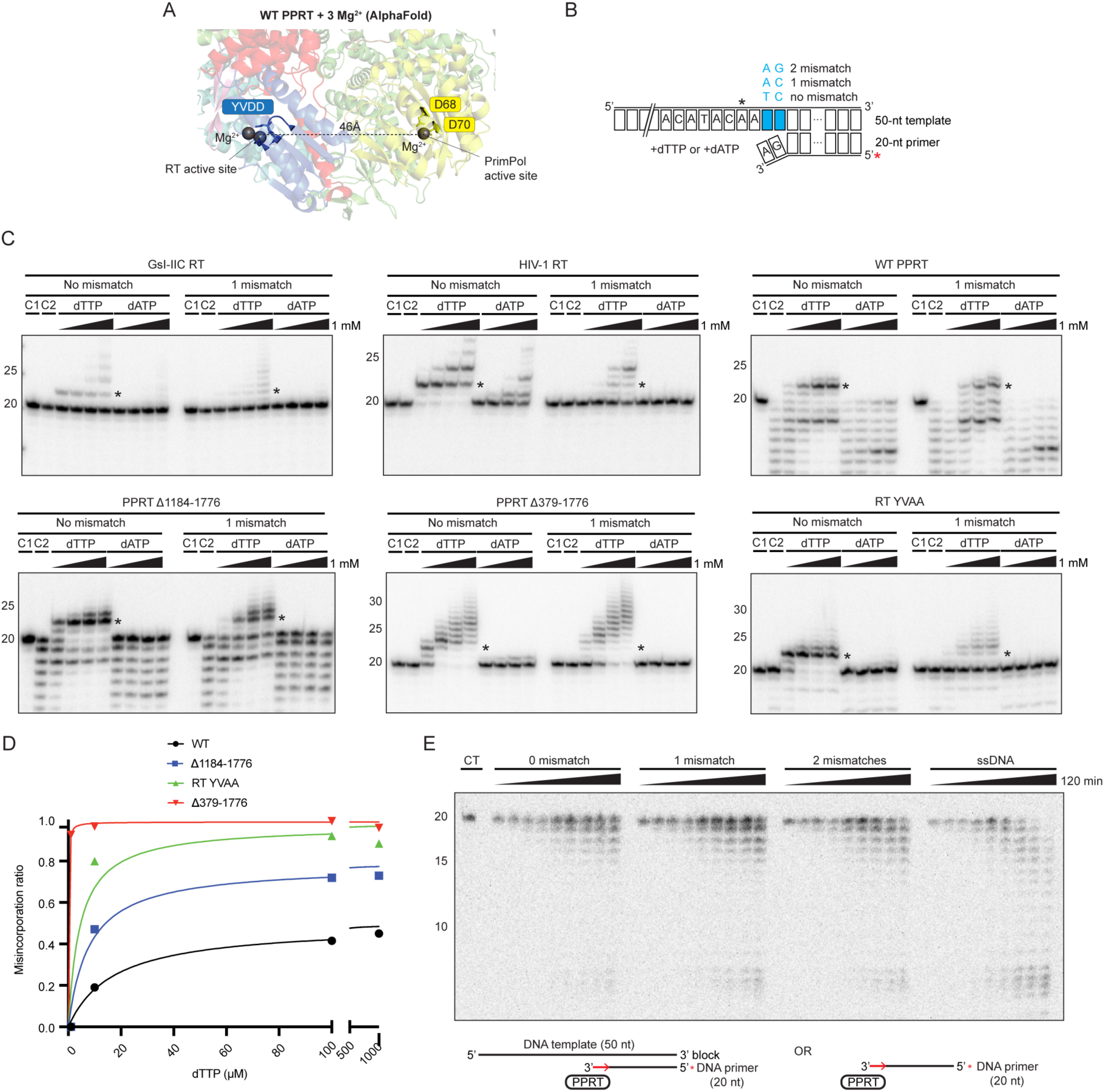
PPRT exonuclease activity functions in proofreading. (A) AlphaFold model of WT PPRT with bound Mg^2+^ ions at the RT (YVDD) and PrimPol (D68 and D70) active sites. (B) Schematic of primer extension substrates comprised of a 3’-Inv(dT)-blocked 50-nt DNA template and 5’-^32^P-labeled (red asterisk) 20-nt DNA primer with fully matched or one or two mismatched template residues opposite the 3’ end of the DNA primer in misincorporation assays (see Methods). The black asterisks in the schematic and gels below indicate the 22-nt position beyond which stalling with a complementary primer would occur after addition of dTTP. (C) Misincorporation assays of GsI-IIC RT, HIV-1 RT, WT PPRT and PPRT mutants using a matched or mismatched DNA primer extension substrate (see above) in reaction medium containing 100 nM enzyme, 10 mM NaCl, 10 mM MgCl_2_, 1 mM MnCl_2_, and 20 mM Tris-HCl pH 7.5 at 37°C for 30 min. GsI-IIC RT was assayed in the absence of MnCl_2_ to suppress terminal transferase activity. In control 1 (C1), the substrate was incubated in reaction medium without protein and dNTPs. In control 2 (C2), the substrate was incubated with the protein but without dNTPs. DNA products were analyzed in a denaturing 17% polyacrylamide gel against 5’-labeled oligonucleotide size markers in a parallel lane (see Methods). (D) Misincorporation ratio of dTTP as a function of dTTP concentration (μM) for PPRT WT (black circles), Δ1184-1776 deletion mutant (blue squares), RT YVAA mutant (green triangles), and Δ379-1776 deletion mutant (red inverted triangles). The misincorporation ratio was calculated as the sum of band intensities ≥23 nt divided by the sum of band intensities ≥22 nt. (E) Exonuclease assay of WT PPRT with a 5’-^32^P-labeled (red asterisk) 20-nt DNA primer and 3’-Inv(dT)-blocked 50-nt DNA template with 0, 1, or 2 mismatched nucleotides at the 3’ end of the primer in reaction medium containing 100 nM enzyme, 10 mM NaCl, 10 mM MgCl_2_, and 20 mM Tris-HCl pH 7.5 at 37°C for times up to 120 min. DNA products were analyzed in a denaturing 17% polyacrylamide gel against 5’-labeled oligonucleotide size markers in a parallel lane (see Methods). CT (control) was tested without enzyme using 0 mismatched substrates under identical conditions, with an incubation time of 120 min.

As an initial test, we compared primer extension reactions with bacterial group II intron-encoded GsI-IIC RT and retroviral HIV-1 RT, neither of which have a 3’ to 5’ exonuclease activity (Figure 5C, top left and middle panels). Consistent with previous findings that GsI-IIC RT has higher fidelity than HIV-1 RT^49^, in reactions with dTTP using a fully matched template-primer combination GsI-IIC RT stalled at 22 nt after addition of two complementary dT residues (band indicated by black asterisk), except at the highest tested dTTP concentrations (100 μM and 1 mM), while HIV-1 RT exhibited substantial misincorporation beyond 22 nt even at lower dTTP concentrations. Likewise, for reactions with the same substrate GsI-IIC RT showed minimal primer extension with dATP, whereas HIV-1 RT displayed higher misincorporation with dATP at non-complementary template nucleotides due to its lower fidelity. With a primer having a single mismatched 3’ nucleotide, HIV-1 RT also had a much higher primer extension activity than GsI-IIC RT with dTTP, further reflecting its lower fidelity.

Having validated these assays, we next used them to compare WT and a series of mutant PPRTs that were found above to have decreased exonuclease activity in the order: WT, Δ1184-1776, RT YVAA, and Δ379-1776, which corresponds to the PrimPol domain only (Figures 3C and 3D). In these assays, WT PPRT primer extension stopped at 22 nt after incorporating two complementary dT residues with both matched and mismatched 3’ primer ends, the latter reflecting removal of a non-complementary nucleotide from the 3’ end of the primer, and showed no detectable extension with dATP even at the highest dATP concentrations tested, indicating high fidelity (Figure 5C, top right panel). PPRT Δ1184-1776, the C-terminal domain deletion mutant that retained moderate exonuclease activity (Figure 3E), largely maintained proper stalling at position 22 nt after the addition of two complementary dT residues. but showed moderately increased misincorporation with both matched and mismatched primer ends at high dTTP concentrations (Figure 5C, bottom left panel). The PPRT mutant YVAA, with very low residual exonuclease activity, maintained proper stalling at 22 nt with matched primers ends, but extended mismatched primer ends beyond this point (Figure 5C, bottom right panel), indicating compromised fidelity due to decreased exonuclease activity. By contrast, PPRT Δ379-1776, which completely lacks exonuclease activity, exhibited longer promiscuous extensions beyond 22 nt with dTTP with both matched and mismatched primer ends, indicating much lower fidelity (Figure 5C, bottom middle panel). As expected, misincorporation by WT and mutant PPRTs was increased at higher dNTP concentrations, reflecting competition between primer extension and exonuclease activity (time courses, Figure 5D). Overall, WT PPRT exhibited higher fidelity than WT GsI-IIC RT and HIV-1 RT, and PPRT variants with diminished exonuclease activity showed compromised fidelity during primer extension, demonstrating the critical role of exonuclease activity in PPRT’s proofreading function.

To further assess PPRT proofreading exonuclease activity, we used primer extension substrates comprised of 3’-blocked 50-nt DNA templates and a 5’-labeled 20-nt primer with complementary or singly or doubly mismatched 3’ residues for different templates and compared exonuclease digestion of the annealed primer to that of the single-stranded DNA primer alone (Figure 5E). WT PPRT showed the strongest 3’ to 5’ exonuclease activity with a ssDNA primer, as expected, followed by double-mismatched, single-mismatched, and non-mismatched primers, respectively. These findings reflect that a ssDNA primer by itself directly accesses the exonuclease active site, while mismatches favor fraying of the 3’ end of an annealed DNA primer, facilitating its access to the exonuclease active site. Thin-layer chromatography and gel electrophoresis of DNA products from exonuclease assays using 3’-labeled 20-nt primers (Figures S3A and S3B) demonstrated that PPRT degrades ssDNA substrates to produce dNMPs, indicating that PPRT uses water as the nucleophile in a hydrolysis-based reaction similar to known proofreading DNA polymerases^50^. Exonuclease assays with different donors (ATP, Pi, PPi) indicated that the inorganic phosphates (Pi and PPi) were not preferred nucleophiles for the exonuclease activity of PPRT (Figure S3C). Collectively, these findings indicate that PPRT’s RT-like domain functions as a conventional proofreading exonuclease, increasing the fidelity of DNA synthesis by the PrimPol domain.

### PPRT proofreading activity mitigates misincorporation due to oxidative DNA damage

To investigate how PPRT’s exonuclease activity might contribute to its cellular response to oxidative DNA damage, we did primer extension assays with a 5’-labeled 20-nt DNA primer and 3’-blocked 50-nt DNA templates that have either an unmodified dG residue, an 8-oxo-dG, or an apurinic/apyrimidinic (AP) site located 23 nt from its 3’-end (asterisk in Figures 6A, 6B, and 6C). These assays showed that WT PPRT, which has both primer extension and DNA exonuclease activities, could readily read through an 8-oxo-dG lesion as efficiently as through an unmodified dG residue at this position (Figure 6B). The YVAA mutant protein, which retains PrimPol primer extension activity, but has minimal RT-like domain exonuclease activity, bypassed the 8-oxo-dG lesion even more efficiently than the WT enzyme, enabling synthesis of a full-length DNA product (Figure 6B). Neither WT nor the mutant enzyme was able to efficiently bypass an AP site at the same position (Figure 6B), stalling mostly at 22 nt.

**Figure 6.**
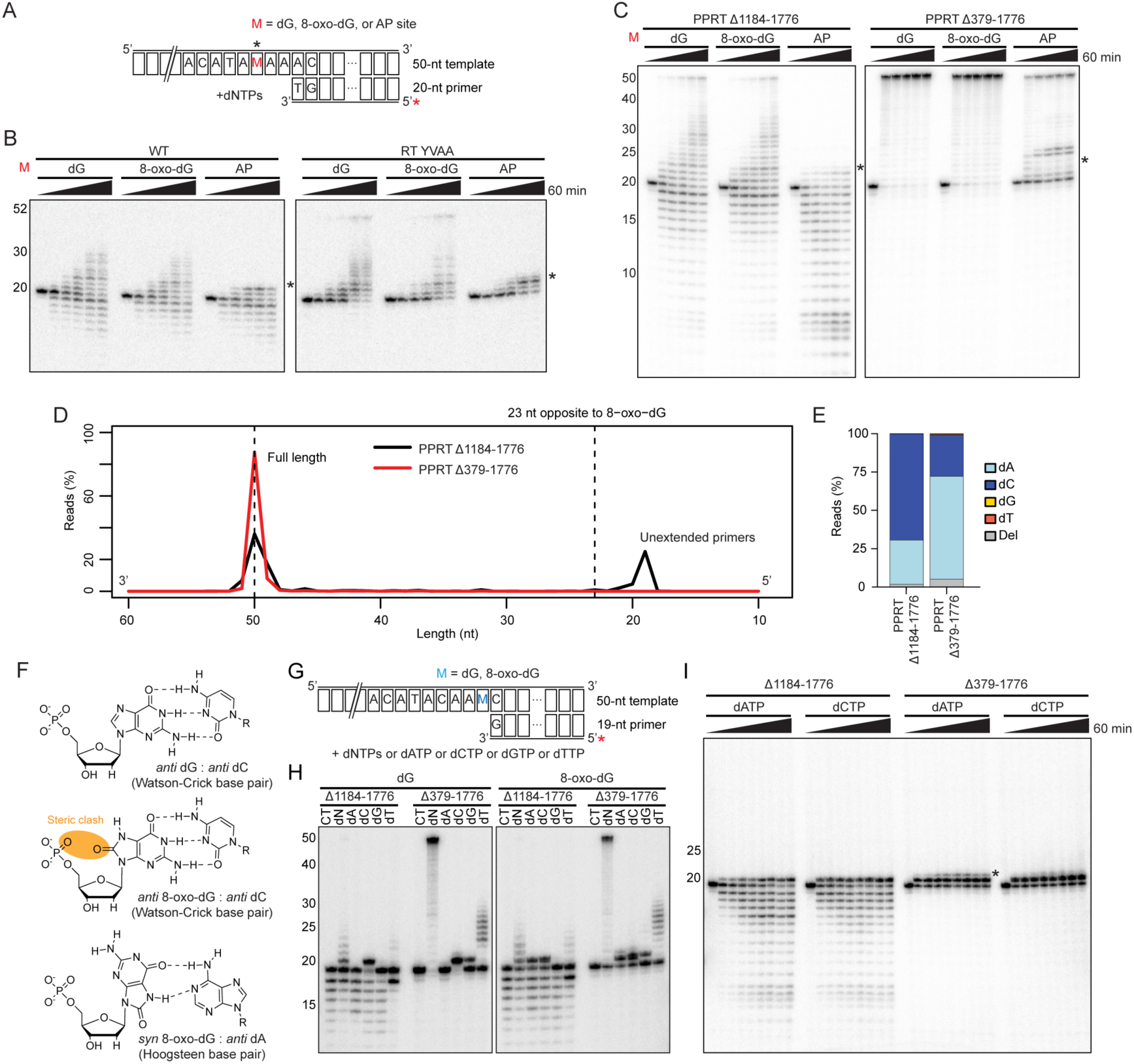
PPRT has 8-oxo-dG translesion DNA synthesis activity. (A) Schematic of DNA damaged substrate used in translesion DNA synthesis assays (see Methods). The substrate is a 3’-Inv(dT)-blocked 50-nt DNA template containing either dG, 8-oxo-dG, or an apurinic/apyrimidinic (AP) site 23 nt from its 3’ end with an annealed 5’-^32^P-labeled (red asterisk) 20-nt DNA primer. Black asterisks in the schematic and gels in panel B indicate the position at which stalling would occur due to a lesion 23 nt from 3’ end of the template. (B) Translesion primer extension assays of WT and YVAA mutant PPRT proteins in reaction medium containing 100 nM enzyme, 10 mM NaCl, 10 mM MgCl_2_, 1 mM MnCl_2_, 20 mM Tris-HCl pH 7.5, and 1 mM dNTPs at 37°C for times up to 60 min. The DNA products were analyzed in a denaturing 17% polyacrylamide gel against 5’-labeled oligonucleotide size markers in a parallel lane (see Methods). (C) Translesion DNA synthesis assays of PPRT C-terminal deletion mutants Δ1184-1776, and PPRT Δ379-1776 done as in panel B. (D) Distribution of MiSeq sequencing read lengths (nt) for DNA products from translesion assays using 1 mM dNTPs for PPRT Δ1184-1776 (black line) and PPRT Δ379-1776 (red line). The vertical dashed line at 23 nt indicates the position opposite to the 8-oxo-dG lesion. The peak at ~20 nt for PPRT Δ1184-1776 corresponds to unused primers, while the peak at ~50 nt corresponds to full-length primer extension products. (E) Stacked bar graph based on the Illumina MiSeq sequencing reads showing the percentage of each nucleotide residue (dA, dC, dG, dT) incorporated at the position opposite the 8-oxo-dG lesion for PPRT C-terminal deletion mutants Δ1184-1776 (left bar), which retains both primer extension and exonuclease activities and PPRT Δ379-1776 (right bar), which retains only the PrimPol domain primer extension activity. Each color represents a different nucleotide residue as indicated in the Figure. The data represent the percentage of total reads containing each nucleotide at this position. (F) Three possible nucleotide base pairings involving a guanine (G) residue or an 8-oxo-dG lesion with cytosine (C) or adenine (A). The top panel shows the canonical Watson-Crick base pair between dG*_anti_* and dC*_anti_*. The middle panel shows a modified base pair involving 8-oxo-dG*_anti_* paired with dC*_anti_*, where the oxidative modification at the C8 position of guanine would cause a steric clash (orange) with the phosphate backbone. The bottom panel shows 8-oxo-dG*_syn_* paired with dA*_anti_* in Hoogsteen geometry, representing a mutagenic mismatch that can trigger a C to A mutation. Hydrogen bonds are depicted as dashed lines, and R represents sugar-phosphate backbone connections. *Anti* and *syn* describe how a nucleobase is oriented relative to its sugar. In the *anti* conformation, the base points away from the sugar; in the *syn* conformation, the base points toward the sugar. (G) Schematic of DNA damaged substrate used in 8-oxo-dG fidelity assays (see Methods). The substrate is a 3’-Inv(dT)-blocked 50-nt DNA template containing an unmodified dG residue or an 8-oxo-dG lesion positioned 20 nt from the 3’-end of an annealed 5’-^32^P-labeled (red asterisk) 19-nt DNA primer. (H) 8-oxo-dG fidelity assay of PPRT Δ1184-1776, which retains exonuclease activity, and PPRT Δ379-1776 (PrimPol domain only), which lacks exonuclease activity, done as in panel B with 1 mM of all or each dNTP as indicated above the autoradiogram at 37°C for 10 min. CT (control) was tested without dNTPs under identical conditions, with an incubation time of 10 min. (I) 8-oxo-dG fidelity assay of PPRT Δ1184-1776 and PPRT Δ379-1776 done as in panel B with 1 mM dATP or dCTP at 37°C for times up to 60 min. The black asterisk in the gel indicates an additional 21-nt band found in the presence of dATP.

We similarly tested two C-terminal deletion mutants: PPRT Δ1184-1776, which retains the PrimPol domain primer extension activity and moderate RT-like domain exonuclease activity without its competing protein-primed DNA synthesis activity, and PPRT Δ379-1776, which retains only the PrimPol domain primer extension activity and lacks both of the other activities. Both mutants bypassed an 8-oxo-dG site without significant stalling at 22 nt (Figure 6C). PPRT Δ1184-1776 showed significant stalling at the AP site at 22 nt, while PPRT Δ379-1776 demonstrated translesion activity at the AP site, synthesizing full-length DNA products but with some products ending at upstream positions 24-25 nt (Figure 6C).

We sequenced the primer-extended products from PPRT Δ1184-1776 and PPRT Δ379-1776 to determine whether PPRT exonuclease activity influences misincorporation during 8-oxo-dG lesion bypass. Both proteins successfully bypassed the 8-oxo-dG lesion at position 23, producing 50-nt full-length cDNA products, although PPRT Δ1184-1776, which retains exonuclease activity, did so with lower efficiency (Figure 6D). DNA sequencing revealed distinct incorporation patterns between the two variants (Figure 6E). PPRT Δ379-1776, lacking proofreading activity, showed a 27:67% dC:dA incorporation ratio opposite the 8-oxo-dG site with 5% deletions at this position. By contrast, PPRT Δ1184-1776, which possesses moderate proofreading activity, preferentially incorporated the complementary dC over dA at a 69:29% ratio and displayed fewer deletions (2%) at the 8-oxo-dG site.

Given that 8-oxo-dG base pairing typically causes a C to A mutation through Hoogsteen base pair formation (Figure 6F)^51–53^, we performed fidelity assays using 5’-labeled 19-nt DNA primers and 3’-blocked 50-nt DNA templates with either an unmodified dG residue or an 8-oxo-dG lesion that is positioned 20 nt from the 3’-end of the template (Figure 6G). The assays were done in the presence of 1 mM of all four or each individual dNTP (denoted dA, dC, dG, or dT in Figure 6H). Both Δ1184-1776 and Δ379-1776 elongated the primer when all four dNTPs were present but with elongation much more efficient for Δ379-1776 due to its lack of exonuclease activity (Figure 6H). However, single dNTP incorporation revealed different fidelity profiles in agreement with Figure 6E: Δ1184-1776, which retains exonuclease, showed a preference for incorporating a dC rather than a dA, while Δ379-1776 incorporated each single dNTP (Figure 6H, right panel). The dT incorporation by Δ379-1776 likely reflects slippage mutations caused by two consecutive dA residues upstream of the modification site. Time-course 8-oxo-dG fidelity assays showed that Δ1184-1776 reached a plateau more slowly with dATP than with dCTP, suggesting that 8-oxo-dG:dA mispairs were more susceptible to exonuclease proofreading activity than 8-oxo-dG:dC pairs and explaining why dC is favored over dA in Figure 6E. By contrast, Δ379-1776 showed no significant difference in incorporation kinetics between dATP and dCTP, but with dATP resulting in an additional 21-nt band (asterisk) that was also minimized for Δ1184-1776 (Figure 6I). Collectively, these findings indicated that PPRT could efficiently read through 8-oxo-dG DNA damage with its proofreading exonuclease activity decreasing misincorporation at this location and at an upstream position due to slippage, beneficial attributes for an enzyme involved in cellular response to oxidative DNA damage.

### Transfer of PPRT’s proofreading exonuclease activity to a group II intron and retroviral RT

The AlphaFold 3.0 predicted structure of the *E. coli* PPRT protein showed that the N-terminal region of the RT domain, which encompasses the fingers and palm regions responsible for template and dNTP binding and includes the RT active site in RT5, is structurally similar to that of GsI-IIC RT, but with a distinctive sequence and predicted structural differences in the RT6/7 region (Figure 7A). In other RTs, RT7 includes the primer grip, which positions and orients the 3’-OH end of the primer strand at the active site, impacting the fidelity of DNA synthesis (Figure 7A)^54^. Inspired by DNA-dependent RNA polymerases, which evolved to integrate polymerase and proofreading activities at the same active site^55^, we aimed to transfer the proofreading exonuclease activity of PPRT to group II intron GsI-IIC and retrovirus HIV-1 RTs by creating hybrid enzymes that fuse their N-terminal region, which contains the RT fingers and palm regions and the RT active site, to PPRT RT6/7 and full-length CTD, which enables maximum DNA exonuclease activity (Figures 3D, 3E, and 7B). GsI-IIC RT was chosen because of its structural similarities to the PPRT RT domain (Figure 1G) and its high thermostability, fidelity, and processivity, which enable efficient synthesis of long primer extension products^56^, and HIV-1 RT was chosen as an extensively characterized retroviral enzyme with high catalytic activity but whose high mutation rate limited its biotechnological applications^48^.

**Figure 7.**
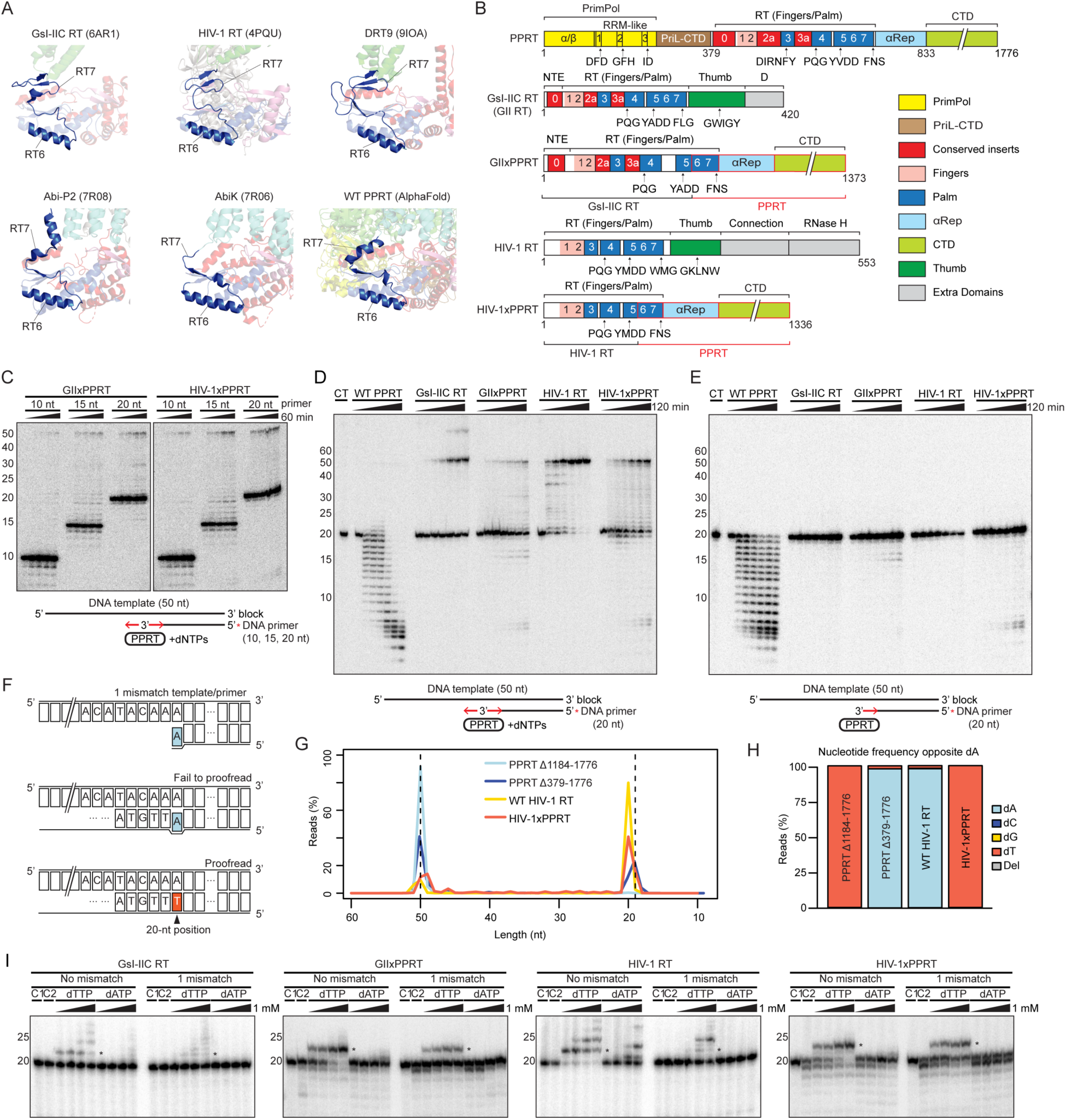
Hybrid proteins comprised of N-terminal RT1-5 active site regions of GsI-IIC or HIV-RT fused to RT6/7 and the C-terminal domain of PPRT enable proofreading at the GsI-IIC and HIV-1 RT active sites. (A) AlphaFold 3 model of PPRT compared with GsI-IIC RT, HIV-1 RT, DRT9, Abi-P2, AbiK, and WT PPRT highlighting the determined or predicted secondary structure RT6 and RT7 motifs of each RT. (B) Schematic of *E. coli* SC366 PPRT, GsI-IIC (GII), and HIV-1 RT and hybrid proteins comprised of N-terminal regions of GII or HIV-1 RT fused to RT6/7 and full-length CTDs of PPRT (denoted GIIxPPRT and HIV-1xPPRT). Protein regions are color-coded as indicated to the right. (C) Primer extension assays of GIIxPPRT and HIV-1xPPRT hybrids using a 3’-Inv(dT)-blocked 50-nt DNA template and 5’-^32^P-labeled (red asterisk) 10, 15, or 20-nt DNA primers in reaction medium containing 100 nM enzyme, 10 mM NaCl, 10 mM MgCl_2_, 20 mM Tris-HCl pH 7.5, and 1 mM dNTPs at 37°C for times up to 120 min. The DNA products were analyzed in a denaturing 17% polyacrylamide gel against 5’-labeled oligonucleotide size markers in a parallel lane (see Methods). (D) Primer extension assays of WT PPRT, WT GsI-IIC RT, GIIxPPRT, WT HIV-1 RT, and HIV-1xPPRT were done as in Panel C using a 3’-Inv(dT)-blocked 50-nt DNA template and a 5’-^32^P-labeled (red asterisk) 20-nt DNA primer at 37°C for times up to 120 min. CT (control) was tested without enzyme under identical conditions, with an incubation time of 120 min. (E) Exonuclease assays of WT PPRT, WT GsI-IIC RT, GIIxPPRT, WT HIV-1 RT, and HIV-1xPPRT done as in panel D in the absence of dNTPs. CT (control) was tested without enzyme under identical conditions, with an incubation time of 120 min. (F) Schematics of template-primer substrates containing a mismatch at the 3’ nucleotide (position 20) of the DNA primer. The diagram shows two possible outcomes for the primer-extension products: failure to proofread resulting in a mismatched A residue at the 20-nt position (light blue), or successful proofreading resulting in a complementary T residue at the 20-nt position (orange). (G) Length distribution (nt) of primer extension products of four polymerases with primers having a mismatched non-complementary 3’ nucleotide determined by Illumina MiSeq sequencing: PPRT Δ1184-1776 (light blue line), PPRT Δ379-1776 (PrimPol domain only; dark blue line), WT HIV-1 RT (yellow line), and HIV-1xPPRT (orange line). The analysis includes both extended and unextended primers. The two vertical dashed lines indicate the ~20 nt mismatched position and the ~50 nt full-length DNA. (H) Nucleotide incorporation frequency at the 20-nt position for extended DNA products analyzed by Illumina MiSeq sequencing using the four different polymerases in panel G. Each stacked bar graph shows the percentages of reads containing different nucleotides at the mismatched 20-nt position color-coded as indicated in the Figure. (I) Misincorporation assays of GIIxPPRT, HIV-1xPPRT, GsI-IIC RT, and HIV-1 RT using a matched or mismatched DNA primer extension substrate (see Methods) in the same reaction medium as panel C with increasing concentrations of dTTP or dATP at 37°C for 30 min. The DNA products were analyzed in a denaturing 17% polyacrylamide gel against 5’-labeled oligonucleotide size markers in a parallel lane (see Methods). In control 1 (C1), the substrate was incubated in reaction medium without protein and dNTPs. In control 2 (C2), the substrate was incubated with the protein but without dNTPs. The black asterisks in the gels indicate the 22-nt position beyond which stalling with a complementary primer should occur after addition of dTTP.

Primer extension assays were performed using purified GIIxPPRT and HIV-1xPPRT enzymes with a 3’-blocked 50-nt DNA template and 5’-labeled 20-nt DNA primer in the presence of 1 mM dNTPs. Optimization of reaction conditions revealed that both of these hybrid RT proteins exhibited maximal primer extension activity at 10 mM Mg^2+^ (Figure S4A), while the addition of 1 mM Mn^2+^ decreased primer extension activity compared to reactions containing only Mg^2+^ (Figure S4B). Testing primers of varying lengths (10, 15, and 20 nt) showed that while primer extension activity was favored with longer primers, exonuclease activity was favored with shorter primers, an inverse relationship, reflecting competition between these activities (Figure 7C). Both the GIIxPPRT and HIV-1xPPRT hybrid proteins exhibited exonuclease activity that digested the 3’ end of an annealed primer in the presence or absence of dNTPs, while no such activity was observed for WT GsI-IIC RT and HIV-1 RT (Figures 7D and 7E). Additionally, both fusion proteins had higher primer extension activity than WT PPRT (Figure 7D), indicating that the efficient primer extension activity of the parental GsI-IIC RT and HIV-1 RTs was successfully incorporated into the engineered enzymes.

To assess the proofreading ability of these hybrid proteins, we compared their primer extension activity with a substrate having a mismatched dA residue at the 3’ end of a 20-nt primer (Figure 7F) to PPRT proteins with shorter (Δ1184-1776) or longer C-terminal deletions (Δ379-1776) that respectively retain or lack the 3’ to 5’-exonuclease activity. Both of these PPRT C-terminal deletion mutants lacked the RT domain protein-primed DNA synthesis activity, which would compete with primer extension activity (Figure 4C). DNA sequencing of the primer extension products confirmed that all of these proteins synthesized full-length extended DNA products (Figure 7G). PPRT Δ379-1776, which corresponds solely to the error-prone PrimPol domain, retained the mismatched dA in nearly all extended products, as did HIV-1 RT, which lacks an inherent exonuclease activity (Figures 7H). By contrast, nearly all of the cDNAs synthesized by the PPRT Δ1184-1776 mutant, which retains exonuclease activity, and the HIV-1xPPRT hybrid protein, which acquired exonuclease activity, had a complementary dT residue at the 20-nt position, reflecting removal of the mismatched dA residues at the 3’ end of the primer and enabling incorporation of a complementary dT residue by primer extension (Figures 7H). To further assess fidelity of the hybrid proteins, we used gel-based misincorporation assays of primer extension with a complementary versus non-complementary dNTP done as in Figure 5B and 5C. These assays showed that both GIIxPPRT and HIV-1xPPRT had higher fidelity than WT GII or HIV-1 RT, as judged by more efficient and discrete extension with a complementary dTTP (asterisk) and minimal mismatched extension with a non-complementary dATP, using either a complementary or mismatched template-primer substrate even at highest dNTP concentration (1 mM; Figure 7I). Collectively, these findings demonstrate a method by which PPRT’s proofreading exonuclease activity can be incorporated into other RTs.

## Discussion

Here, we found that an *E. coli* PPRT protein (DRT7, UG10), which functions in phage defense^17^, has an additional cellular function in repair of oxidative DNA damage as well as an unprecedented 3’ to 5’ exonuclease activity that is associated with the RT-like domain and functions in proofreading DNAs synthesized by the error-prone PrimPol domain. We also found that the long (~1,000 amino acid) C-terminal region (αRep plus the CTD) of the PPRT protein contributes to these activities, likely by stabilizing the structure or relative positions of the PrimPol and RT-like domains and/or providing enclosed channels for their biochemical activities. Extending these findings, we identified predicted structural features of the RT-like domain of the PPRT protein that contribute to its proofreading exonuclease activity, enabling us to engineer this activity into hybrid proteins comprised of an N-terminal group II intron or retrovirus RT domain containing the RT active site fused to the altered RT6/7 primer grip region and the C-terminal domain of the PPRT protein needed for maximum DNA exonuclease activity. These findings demonstrate one method and suggested additional methods (see below) for engineering a proofreading activity into RTs, enabling the development of higher fidelity enzymes for use in biotechnological and gene therapy applications.

Our initial biochemical analysis revealed that bacterial PPRT’s PrimPol domain has functional similarities to a human CCDC11 PrimPol protein, particularly in DNA damage tolerance and replication stress response^43^. While other bacterial proteins with a PrimPol domain have been found to act as barriers against naturally acquired DNA^57^ and participate in CRISPR-Cas spacer acquisition^58^, PPRT’s PrimPol domain has biochemical characteristics that closely mirror those of human PrimPol proteins. These include a preference for Mn^2+^ ions for DNA synthesis (Figure 2A); a weak ability to bypass an abasic site (Figures 6B and 6C); and the ability to efficiently read through 8-oxo-dG lesions (Figures 6B and 6D), which contributes to a cellular response to oxidative DNA damage (Figure 1J)^43,59–62^.

A striking difference between the PPRT and human PrimPol proteins, however, lies in the error prevention mechanisms of these enzymes. Human PrimPol enzymes lack a 3’ to 5’ exonuclease activity^43^ and are considered error-prone^59,61,63^. Surprisingly, while many DNA polymerase families have a separate evolutionarily conserved proofreading exonuclease domain with consensus motifs (ExoI, ExoII, and ExoIII)^64,65^, we found that the fused RT-like domain of PPRT possesses an inherent 3’ to 5’ exonuclease activity that is dependent upon the YVDD motif that binds catalytic divalent cations at the RT active site and enables proofreading of DNAs synthesized by the PrimPol domain (Figures 3, 5, and 7). PPRT’s exonuclease activity also produces monophosphates (Figure S3B), which are byproducts of the proofreading exonuclease activity found in DNA polymerases^50^. This unique interacting architecture enables PPRT to maintain high fidelity while performing more accurate translesion DNA synthesis at 8-oxo-dG residues, a beneficial attribute of this sophisticatedly evolved protein.

While a hydrolysis-based 3’ to 5’ exonuclease activity observed in PPRT is a function typically found in the separate exonuclease domain of DNA polymerases^66^ or the active site of DNA-dependent RNA polymerases (RNAPs)^67^, the identification of an RT-like domain that has an inherent proofreading 3’ to 5’ exonuclease activity is unprecedented. Although both the *E. coli* PPRT protein and *Caloramator australicus Ca*CART-CAPP have highly conserved PrimPol and RT domains, the order of these fused domains differs and the N-terminal RT domain of *Ca*CART-CAPP is structurally similar to GsI-IIC RT with a canonical RT activity that functions in RNA spacer acquisition^58^. Comparison of the AlphaFold predicted structure of the RT-like domain of PPRT with the structure of GsI-IIC RT^4^ suggested that PPRT’s exonuclease activity at the RT active site is likely due at least in part to structural differences in RT6/7 (Figures 1F, 4A, and 7A), which lacks a primer grip structure that delivers the 3’ OH end of the primer to the RT active site. The αRep and CTD domains of PPRT most likely function as auxiliary regions that structurally support exonuclease activity at the RT active site rather than contributing directly to this activity.

Abi polymerases (*e.g.*, AbiA, AbiK, and Abi-P2), whose RT domains function in phage defense by ssDNA synthesis, possess conserved RT motifs RT0-6 but lack the RT7 primer grip region with RT0-6 fused directly to C-terminal α-helical domain^17,24,25^. It was previously hypothesized that this α-helical domain may contribute to the template-independent single-stranded DNA synthesis activity of Abi polymerases^24,25^. While PPRT also possesses a similarly positioned α-helical domain, additional sequences of the CTD beyond this α-helical region were essential for optimal ssDNA synthesis and exonuclease activity by the RT-like domain (Figures 3D, 3E, and 4C), possibly by structural interaction and/or constraining templates and extended products. Biochemical analysis indicated that the proofreading activity of PPRT depends upon shifting of the 3’ end of a mismatched primer or nascent DNA between the PrimPol and RT-like domains, similar to the mechanism by which DNA polymerases that have a separate 3’ to 5’ exonuclease act in proofreading^68,69^.

Biochemical analysis also showed that the RT-like domain of PPRT performs protein-primed single-stranded DNA synthesis similar to other DRTs^26,29^, but differs in having a strong preference for synthesizing poly(dT) rather than poly(dA) (Figures 4E, 4F, 4G, and 4H). Remarkably, the initial protein-primed ssDNA product switches between the RT-like and PrimPol domains, and when associated with the latter, generates DNAs with double-stranded regions containing poly(dA) runs (Figures 4D and 4F), a process entirely dependent on the initial poly(dT)-rich DNA product synthesized by the RT domain (Figure 4B). DRT9 RTs, which have been shown to rely upon poly(dA) synthesis for phage defense, differ from PPRT in synthesizing poly(dA) directly via their RT domain using a short poly(U) sequence located in neighboring ncRNA as the template^26,29^. It will be of interest to investigate if the synthesis of partially dsDNA products by PPRT contributes to phage defense via greater dNTP depletion or by other mechanisms.

An initial gene disruption experiment indicated that the WT PPRT, in addition to its previously identified function in phage defense, contributes to a cellular response to oxidative DNA damage. In accord with this finding, biochemical assays showed that the PrimPol domain primer extension activity could efficiently read through 8-oxo-dG, a major contributor to lethal oxidative DNA damage by triggering stalling of replication machinery and C to A mutations^51,52^. Additionally, PPRT’s RT-like domain proofreading exonuclease activity enables read through of 8-oxo-dG at a lower mutation frequency than a mutant protein that lacks this activity (Figures 6E, 6H, and 6I). These findings raise the possibility that other bacterial RTs characterized as functioning in phage defense by simple plaque assays without further characterization might also have additional cellular functions.

Finally, our finding that the PPRT PrimPol domain is fused to an RT domain that functions in proofreading a nascent DNA strand that switches up and back between the two separate active sites suggests a general mechanism for increasing the fidelity of PrimPol DNA polymerases. Likewise, our finding that hybrids in which N-terminal regions of group II intron or HIV-1 RT extending through the RT active site were fused to PPRT’s RT6/7 and C-terminal domain regions enabled proofreading suggests a general mechanism for associating proofreading activity directly with the canonical active site of other RTs, although this may require additional structural modifications to further optimize biochemical activities. Finally, our finding that structural differences in PPRT’s RT6/7 motifs are correlated with 3’ to 5’ exonuclease proofreading activity suggests that engineering structural modifications into these and possibly neighboring regions may be an alternative method of engineering proofreading activity directly into other RTs without constructing hybrid proteins. Higher fidelity RTs constructed by these methods would have widespread RNA-sequencing and genome engineering applications.

## Data and code availability

TGIRT-seq datasets for *E. coli* WT and *pprt* disruptant strains have been deposited in the Sequence Read Archive (SRA) under accession number PRJNA1252753. *In vitro* experimental data and genomic sequences of *E. coli* WT and *pprt* disruptant strains have been deposited in the SRA under accession number PRJNA1280996. Whole genome shotgun sequencing data is available under NCBI accession number NZ_QOMS01000055.1. Code for data analysis has been deposited in Zenodo: DOI:10.5281/zenodo.15722935.

## Acknowledgements

We thank Laura Markham for help with protein preparations. This research was supported by Welch Foundation grant F1607 and NIH grant R35 GM136216 to AML and NSFC (No. 32160015) to GC.

## Author contributions

SK developed an improved PPRT protein purification method, designed and did all biochemical studies and structural modeling in the manuscript, and wrote the manuscript with AML. HX designed all next generation sequencing related methods, performed gene disruption analysis, and wrote the *pprt* gene disruption section. GM initiated the project with HX and GC, analyzed the genomic context and distribution of genes encoding PPRT proteins in different bacteria, and identified the *E. coli* PPRT protein as the candidate for further investigation; JY did bioinformatic analysis of PPRT protein ssDNA synthesis and proofreading DNA exonuclease activities; JYZ, a co-advisor of SK, made suggestions for structural predictions and experiments; GC, a visiting faculty member, initiated the project at the University of Texas at Austin and did gene disruption and initial biochemical analysis that was interrupted by the pandemic; AML worked with other authors on designing experiments and data analysis, oversaw the project, and wrote the manuscript with SK.

## Methods

### AlphaFold structural modeling and Foldseek analysis

Protein sequences were submitted to the AlphaFold 3 webserver^30^, and structural predictions were visualized by using PyMOL^33^. Structural models were submitted to the Foldseek Search Server^31^ to identify structurally similar proteins.

### Bacterial strains

The bacterial strains used in this study are listed in Table S2. *Escherichia coli* SC366 was obtained from Dr. Michael J. Sadowsky (Biotechnology Institute, University of Minnesota, St. Paul, Minnesota, USA). Rosetta 2 (DE3) was purchased from Novagen; Top10 and DH5α were purchased from ThermoFisher (Table S2).

### Plasmids

Plasmids and primers used in the study are listed in Tables S3 and S4 respectively. The pMal-PPRT expression vector, designed to produce PPRT protein fused to an N-terminal Maltose-Binding Protein (MBP) tag via a rigid linker^56^ and with a C-terminal 7xHis tag, was constructed in two steps. In the first step, a 3.9-kb 3’ PPRT segment was amplified from the *E. coli* 366 genome by PCR with primers PPRT-DP1 and PPRT-DP2, digested with *Nco*I and *Pst*I, and then ligated into the *Nco*I and *Pst*I-digested pMal-c5X expression plasmid (New England Biolabs). In the second step, a 1.5-kb 5’ PPRT segment was amplified from the *E. coli* 366 genome by PCR using primers PPRT-UP1 and PPRT-UP2, digested with *Nco*I, and then ligated into the above intermediate plasmid digested with *Nco*I to obtain the expression plasmid pMal-PPRT. The sequence of pMal-PPRT was verified by DNA sequencing (Plasmidsaurus, Eton Bioscience).

pMal-PPRT-D68A and pMal-PPRT-D70A were constructed by using Primer sets PrmS-F/PrmS-R to amplify a 0.4-kb segment of the PrimPol domain, which was digested with *Bam*HI and *Sph*I and ligated into a pCR^TM^2.1 T vector (ThermoFisher) digested with the same enzymes. The resulting plasmid was then used as a template to introduce the point mutations D68A and D70A with primer sets D68A-F/D68A-R and D70A-F/D70A-R, respectively. The final products were digested with *Dpn*I and transformed into *E. coli* strain Top10 (ThermoFisher). After sequencing, the pCR^TM^2.1 T vectors containing the PrimPol domain point mutations D68A or D70A were digested with *Bam*HI and *Sph*I, generating a 0.4-kb DNA fragment that was then ligated into the pMal-PPRT plasmid digested with the same enzymes to construct the PrimPol domain point mutation expression plasmids pMal-PPRT-D68A and pMal-PPRT-D70A.

pMal-PPRT-D68A+D70A was constructed by site-directed mutagenesis of the pMal-PPRT-D68A plasmid using the primer set D70A_2-F/D70A_2-R.

pMal-PPRT-YVAA was constructed by using WT pMal-PPRT as a template with primer set 0104-1/0104-2 and 0104-3/0104-4 to amplify 6.3- and 5.6-kb DNA fragments separately. These two segments were then digested with *Snb*I and *Pci*I and ligated together to construct the YVAA mutant expression plasmid pMal-PPRT-YVAA.

pMal-PPRT-D526A+YVAA was constructed by site-directed mutagenesis of pMal-PPRT-YVAA using primer sets D526A_F/D526A_R.

pMal-PPRT-D68A+D70A+YVAA was constructed by site-directed mutagenesis of pMal-PPRT-YVAA using primer set D68A_2-F/D68A_2-R, followed by a second round of site-directed mutagenesis using primer set D70A_2-F/D70A_2-R.

pMal-PPRTΔ1184-1776 was constructed by digesting pMal-PPRT with *Eco*RI and *Pst*I to yield a 10.1-kb DNA fragment. Primer set 1184-F/1184-R was annealed separately to form a double-stranded DNA linker, which was then digested with the same *Eco*RI and *Pst*I enzymes, and these two segments were ligated with T4 DNA ligase (New England Biolabs).

pMal-PPRTΔ836-1776 was constructed by PCR of WT pMal-PPRT with primer set 836-F/836-R to yield a 9,099-bp DNA fragment, which was then self-ligated using kinase, ligase, and DpnI (KLD) enzyme mix (New England Biolabs).

pMal-PPRTΔ678-1776 was constructed by PCR of pMal-PPRT using primer set 678-F/678-R to yield a 8,025-bp DNA fragment that was self-ligated with KLD enzyme mix (New England Biolabs).

pMal-PPRTΔ379-1776 was constructed by PCR of pMal-PPRT using primer set 379-F/379-R to yield a 7,731-bp DNA fragment that was self-ligated with KLD enzyme mix (New England Biolabs).

pMal-GIIxPPRT, which is comprised of N-terminus of GsI-IIC RT containing RT0 to RT5 fused RT6/7 and the CTD of PPRT, was constructed by PCR of pMal-GII RT^56^ using primer set En_G2-F/En_G2-R to yield a 1,841-bp DNA fragment and PCR of pMal-PPRT with primer set En PPRT-F/En_PPRT-R to yield an 8,912-bp DNA fragment. The two fragments were then assembled using NEBuilder HiFi DNA assembly Master Mix (New England Biolabs).

pMal-HIV-1xPPRT, which is comprised of the N-terminus of HIV-1 RT containing RT1 to 5 fused to RT6/7 and the CTD of PPRT, was synthesized by GenScript.

All plasmids were sequenced to confirm the expected sequence and the absence of secondary mutations.

### Targetron disruption analysis of the *pprt* gene function

The targetron expression plasmid pBL1-PPRT::1218s used for disruption of the *pprt* gene is a derivative of pBL1^70^, which was derived from broad host range expression vector pJB866^71^. The targetron plasmid expresses the Ll.LtrB-ΔORF targetron using an m-toluic acid-inducible promoter and carries a *Tet^R^* marker^70^.

A targetron that inserts at the desired site was designed based on an online tool (https://sites.cns.utexas.edu/lambowitz/targetron-design^70^) and constructed by successive overlap extension PCRs with primer sets 1218IBS/EBSU, 1218EBS2/EBS1d, and 1218IBS/EBS1d (Table S5), a method described previously for programming base-pairing interactions for targetron insertion at desired sites^72^. The final 350-bp PCR product was then digested with *Hind*III and *Bsr*GI and ligated between the same sites in targetron expression plasmid pBL1. The programmed targetron inserts after PPRT nucleotide 1218s (*E. coli* genome position 105767/226080 bp) in the top (sense) strand. Targetron insertion at the desired site was confirmed by Sanger sequencing (Plasmidsaurus, Eton Bioscience).

Transformation and screening of *E. coli* SC366 for *pprt* gene disruption were done as follows: (i) *E. coli* SC366 cells were grown in 50-ml Luria-Bertani (LB) medium at 37°C to early exponential phase (O.D._600_ = 0.4-0.6) and chilled on ice for at least 30 min before harvesting. The harvested cells were then washed twice with ice-cold distilled water, pelleted, and resuspended in 5-ml of ice-cold 10% glycerol, and kept on ice for at least 30 min; (ii) 2 μg of the targetron expression plasmid pBL1-PPRT::1218s was mixed with 100 μL of *E. coli* SC366 competent cells in LB medium containing tetracycline (25 μg/mL), and the mixture was then transferred to a 0.2-cm cuvette (Bio-Rad) and kept on ice for 30 min prior to electroporation. (iii) Electroporation was carried out using Bio-Rad Gene Pulser Xcell system under the following conditions: 200 Ω, 25 μF and 2500 V. (iv) After electroporation, 1-ml of LB medium was added, and the cells were allowed to recover in a rotary shaker (220 rpm, 1 h at 37°C). The recovered cells were then plated on LB agar containing tetracycline (25 μg/ml) and incubated overnight at 37°C; (v) tetracycline-resistant colonies were picked and inoculated into a 10-ml conical tube containing 3-ml LB broth supplemented with tetracycline (25 μg/ml), and the cells were grown for 2 h at 37°C prior to inducing targetron expression with 4 mM m-toluic acid for 5 h at 37°C. 50 μL of cells were pipetted and plated on LB agar supplemented with tetracycline (25 μg/ml) and incubated overnight at 37°C; (vi) transformants with the targetron inserted at the 1218s target site were identified by colony PCR^70^ of extracted genomic DNA from *pprt* disruptant with primers 1218-F and 1218-R (Table S5). *pprt* disruptants gave a 1.3-kb fragment while the wild-type cells yielded a 0.4-kb product (Figure S1C).

To cure the targetron expression plasmid, positive transformants were inoculated and cultivated in 100-ml LB medium at 37°C without any antibiotics. After 20 consecutive transfers (twice per day for 10 days), the cells were plated on LB agar and incubated overnight at 37°C. Individual colonies were then picked and resuspended into 100-μL sterile distilled water and then split equally and reinoculated separately into 100-ml LB liquid medium with or without tetracycline (25 μg/ml). Cells that could only grow at 37°C in LB medium without tetracycline but not in LB medium with tetracycline were selected to verify the loss of the targetron expression plasmid.

Targetron insertion at the desired site was tested initially by Southern hybridization of extracted genomic DNA from *pprt* disruptant with a 197-bp targetron probe (Table S6) generated by PCR using primers IntronPbF and IntronPbR (Table S5) and labeled with [γ-^32^P]-dATP (6,000 Ci/mmol, Revvity) using T4 polynucleotide kinase (New England Biolabs). Genomic DNA was isolated from wild-type and mutant strains using Monarch Genomic DNA purification kit (New England Biolabs) according to the manufacturer’s instructions. The genomic DNA was then digested with *Hind*III and *EcoR*I at 37°C for 2 h, run in a 1.0 % agarose gel, and transferred to a Nylon membrane (Hybond-NX, GE Healthcare), which was hybridized with the ^32^P-labeled targetron probe. A disruptant with a single band detected by Southern hybridization at the desired site was chosen for phenotypic and TGIRT-seq analysis.

Growth curves for the WT *E. coli* SC366 and *pprt* disruptant strains were done in 250-ml Erlenmeyer flasks containing 100-ml LB without antibiotics. To obtain these growth curves, single colonies from an LB agar plate without antibiotics were picked and inoculated into 100-ml LB growth medium and grown overnight in a rotary shaker (220 rpm) at 37°C. At different growth intervals, the cell concentration was monitored by measuring the optical density at 600 nm with a UV-Vis spectrophotometer to select time points for TGIRT-seq analysis of gene expression differences between the WT and *pprt* disruptant strains.

### Preparation of *E. coli* WT and *pprt* disruptant cellular RNAs for TGIRT-seq

Wild-type (WT) *E. coli* SC366 and the *pprt* disruptant strain were grown as described above, and 500-μl samples were collected at 2, 5, and 8 h. Total cellular RNA was extracted by using a Monarch total RNA miniprep kit (New England Biolabs) at the indicated time points. For TGIRT-seq library preparation, cellular RNAs (500 ng) were incubated with Baseline-ZERO DNase (Lucigen; 2 units, 30 min at 37°C) to digest DNA followed by rRNA depletion using an Illumina Ribo-Zero Plus rRNA depletion kit. The volume of the rRNA-depleted RNAs was then brought to 100 μl by adding nuclease free water. Clean-up and size-selection were performed using RNAClean XP beads (Beckman Coulter) with a first-round sample to bead volume (v/v) of 1:0.45 to separate short RNAs (<300 nt) from longer RNAs ≥300 nt that remained bound to the magnetic beads. After separation of the beads, the short RNAs were transferred into a new tube with the addition of fresh beads with a sample to bead v/v ratio of 1:1.8. The beads with bound long and short RNAs were then washed twice with 200 μL of freshly made 80% ethanol for 30 sec each and eluted with nuclease free water. The long RNAs (>300 nt) were chemically fragmented to 70-90 nt length by using an NEBNext Magnesium RNA Fragmentation Module (New England Biolabs; 94°C for 5 min) and cleaned up with a Zymo RNA clean and concentrator kit using a modified 8X ethanol protocol (v/v ratio of 1:2:8 for RNA sample/ kit RNA Binding Buffer/100% ethanol) to minimize loss of very small RNAs^40,73^. The chemically fragmented long RNAs were then combined with the short RNAs (<300 nt), and the reconstituted RNA preparation was treated with T4 polynucleotide kinase (Lucigen; 50 U for 30 min at 37°C) to remove 3’ phosphates and 2’,3’-cyclic phosphates, which impede TGIRT template switching, followed by a final clean-up with a Zymo RNA clean and concentrator kit using the modified 8X ethanol protocol above.

### TGIRT-seq library preparation

TGIRT-seq libraries were constructed as described^74,75^ by using TGIRT-template switching from a synthetic RNA template/DNA primer duplex with a single nucleotide 3’ overhang that initiates cDNA synthesis by base pairing to the 3’ nucleotide of the target RNA for 3’ RNAseq adapter addition followed by single-stranded DNA ligation to the 3’ end of the cDNA using Thermostable 5’ AppDNA/RNA Ligase (New England Biolabs) for 5’ RNA-seq adapter addition. The TGIRT-template-switching DNA synthesis reaction was done with 1 μM TGIRT-III (InGex, LLC; currently available from the Lambowitz laboratory) for 15 min at 60°C. The resulting cDNAs were amplified by PCR with primers that add capture sites and indices for Illumina sequence (denaturation 98°C for 5 s, followed by 12 cycles of 98°C for 5 s, 65°C for 10 s, and 72°C for 10 s). The PCR products were purified by using Agencourt AMPure XP beads (Beckman Coulter) and stored at −20°C prior to sequence analysis.

### Thermostable Group II Intron Reverse Transcriptase Sequencing (TGIRT-seq) of protein-coding and non-coding RNAs

For sequence analysis of protein-coding and non-coding RNAs in *E. coli* SC366, TGIRT-seq libraries prepared from *E. coli* SC366 WT and *pprt* disruptant cellular RNAs were sequenced on an Illumina NovaSeq 6000 at the Genome Sequencing and Analysis Facility (GSAF) at the University of Texas at Austin to obtain ~25 million 2 x 75 nt paired-end reads per sample. Illumina TruSeq adapters and PCR primer sequences were trimmed from the reads with Cutadapt v2.8 (sequence quality score cut-off at 20; p-value <0.01)^76^ and reads <15-nt after trimming were discarded. Reads were then mapped by using HISAT2 with default settings to the *E. coli* SC366 genome reference sequence (NZ_QOMS01000055.1). Uniquely mapped reads and the filtered multiply mapped reads were combined and intersected with gene annotations (NZ_QOMS01000055.1).

Principal Component Analysis plots and bar graphs were generated in R^77^ from the raw read count table. Gene-wise differential expression analyses were performed using DESeq2^78^, and the results were compared after the log-fold-change was shrunk (lfcshrink) by the “ashr” method for higher accuracy^79^. For the design matrix, the DESeq2 normalized count was designed with the consideration of the growth time, genotype of the PPRT gene and the interaction of the two factors with the formula: counts ~ growth time + genotype + (growth time:genotype) (Figure S1H). The inclusion of the interaction term was done to minimize background changes due to the growth times and thus emphasizes changes due to PPRT gene disruption at different time point. Gene-set enrichment analysis (GSEA) was performed from the selected candidate genes list by using the ShinyGO v8.0^42^ and the heatmap was generated in R from the returned ShinyGO results.

The bam files were down-sampled to 5 million reads in a random manner by using samtools. The three bam files for each replicate of genotype/growth time combination were merged, sorted and indexed for the IGV. Genomic variation analyses were performed on the RNA-seq reads with bcftools^80^. The same down-sampled and sorted bam files were used as input for the variant calling by the command: bcftools mpileup -Ob -o “output bcf format file” -f “genome reference.fa file” “input bam file”. The summary of the variants was generated by the command: bcftools call -vmO z -o “output file” “input bcf file”.

### PPRT protein expression and purification

Wild-type and mutant PPRT proteins were expressed in *E. coli* Rosetta2 (DE3). Competent cells were transformed with pMal-c5X (New England Biolabs) vectors carrying the *pprt* WT or mutant genes according to the manufacturer’s protocol and selected by plating on agar containing LB medium plus 100 μg/ml ampicillin and 25 μg/ml chloramphenicol at 37°C. A single colony was then inoculated in LB broth supplemented with the same antibiotics and incubated overnight at 37°C in a rotary shaker. 5 ml of the overnight culture was reinoculated into 100-ml fresh LB broth supplemented with the same antibiotics and allowed to grow to log phase (O.D._600_ ≈ 0.7). 10-ml of the 100-ml cultures was then added to 1 liter of fresh LB broth supplemented with the above antibiotics and incubated for 2-3 h in a shaker (220 rpm, 37°C). When the culture reached O.D._600_ ≈ 0.7, isopropyl β-D-1-thiogalactopyranoside was added to a final concentration of 0.5 mM, and the cultures were incubated for 18 h in a shaker at 130 rpm and 18°C. Cells were harvested by centrifugation, and the pellet was resuspended in 35-ml of ice-cold buffer A1 (25 mM Tris-HCl, pH 7.5; 500 mM NaCl). The collected cells were then treated with lysozyme (1 mg/ml, 0.5 h, 4°C) and sonicated (Branson Sonifier 450; 10 bursts of 0.5 min each with 0.5 min between each burst). The supernatant was collected by centrifugation (15,000 x g, 30 min, 4°C) and polyethyleneimine (PEI) was added drop by drop with stirring to a final concentration of 0.2%. The mixture was kept on ice for 10 min to precipitate nucleic acids, and the supernatant was collected by centrifugation in an Avanti J-E centrifuge (Beckman) (15,000 x g, 30 min, 4°C). Ammonium sulfate was then added to the supernatant to a final saturation of 60%. After 30 min on ice, the crude proteins were collected by centrifugation in an Avanti J-E centrifuge (Beckman) (15,000 x g, 30 min, 4°C) and dissolved in 15-ml A1 buffer (25 mM Tris-HCl, pH 7.5; 500 mM NaCl). Prior to purification with an ÅKTA start system (Cytiva), crude proteins were filtered through a 0.45-μm polyethersulfone membrane syringe filter (Pall Corporation) and then loaded onto a 5-ml MBPTrap-HP-column (Cytiva). The column was washed sequentially with buffers A1 (25 mM Tris-HCl pH 7.5, 500 mM NaCl), A2 (25 mM Tris-HCl pH 7.5, 1.5 M NaCl), and A1 again (3-column volumes each). The bound proteins were eluted with buffer A3 (buffer A1 + 10 mM maltose), and the eluted fractions containing PPRT protein were confirmed by SDS-PAGE, pooled and loaded onto a 5-ml Ni-NTA HisTrap HP column (Cytiva). The column was washed with buffer A4 (buffer A1 + 30 mM imidazole) and A5 (buffer A1 + 45 mM imidazole), followed by elution with buffer A6 (buffer A1 + 300 mM imidazole). The purified PPRT proteins were dialyzed overnight against storage buffer (25 mM Tris-HCl pH 7.5, 100 mM NaCl, 50% glycerol), concentrated to 5-10 mg/ml with an Amicon Ultra-15 (100K) or Amicon Ultra-0.5ml (100K) concentrator (Millipore) according to manufacturer’s protocol, and stored at −80°C. The final preparation had an A_260_/A_280_ ratio of 0.6-0.7, indicating minimal nucleic acid contamination.

The GIIxPPRT hybrid protein was purified as described above for PPRT proteins, but the HIV-1xPPRT hybrid protein required a modified purification protocol by skipping the PEI-purification step as the protein was lost during this step. Crude proteins were filtered through a 0.45-μm polyethersulfone membrane syringe filter (Pall Corporation) and then loaded onto a 5-ml MBPTrap-HP-column (Cytiva). The column was washed sequentially with buffers A1, A2, and A1 again (3-column volumes each). The bound proteins were eluted with buffer A3, and the eluted fractions containing PPRT protein were confirmed by SDS-PAGE, then pooled and diluted to 25 mM Tris-HCl pH 7.5, 100 mM NaCl before being loaded onto a 5-ml Hi-Trap Heparin HP column (Cytiva). The column was washed with buffer A4-2 (25 mM Tris-HCl pH 7.5, 100 mM NaCl) and the bound proteins were eluted using 10 column volumes of a 0.1 to 1.5 M NaCl gradient collecting 2-mL fractions. The purified PPRT proteins were dialyzed overnight against storage buffer (25 mM Tris-HCl pH 7.5, 100 mM NaCl, 50% glycerol), concentrated to 1-2 mg/ml, and stored at −80°C. Purified proteins (0.5 μg) used in this study were run on NuPAGE 4-12 % Bis-Tris gel and stained with Coomassie blue (Figure S5)

### Biochemical assays

Unless indicated otherwise, biochemical assays of wild-type and mutant PPRT protein were done at 37°C with purified protein in reaction medium containing 10 mM NaCl, 10 mM MgCl_2_, 1 mM MnCl_2_, and 20 mM Tris-HCl pH 7.5 (denoted reaction medium below). Single reactions were done in 10 μl of reaction medium, and time courses were done in 100 μl of reaction medium with 10 μl taken at each time point. Reactions were stopped by adding 2 μl of STOP solution (250 mM EDTA; Sigma-Aldrich), 400 U/ml proteinase K (New England Biolabs) and incubating at 37°C for 30 min for protein digestion. After adding 6 μl of 3x formamide loading dye, the mixture was denatured by heating to 95°C for 3 min prior to electrophoresis. Products were analyzed in an 8 M urea/17% polyacrylamide gel run at 55 W for 1.5-2 h against 5’-labeled oligonucleotide size markers, either 10 to 60 or 20 to 100 nt DNA oligonucleotide length standards (IDT) in a parallel lane. After drying, the gel was scanned with a phosphorimager (Typhoon FL A 9500, GE Healthcare).

Primer extension assays were carried out by preparing a 20 nM 5’-^32^P-labeled 20-nt DNA primer (3’-TG, Table S4) annealed to a 22 nM 3’-Inv(dT)-blocked 50-nt DNA (5’-AC, Table S4) to prevent snapback DNA synthesis. The 5’ end of the primer was labeled with [γ-^32^P]-ATP (6,000 Ci/mmol; Revvity) using T4 polynucleotide kinase (PNK, New England Biolabs) followed by clean up using an Oligo Clean & Concentrator Kit (Zymo Research). The labeled DNA primer was annealed to the DNA template at a molar ratio of 1:1.1 by heating to 95°C for 3 min followed by slowly cooling to room temperature. 100 nM PPRT was incubated with 20 nM of the annealed template-primer substrate in reaction medium containing 1 mM dNTPs (final concentration 1 mM each of dATP, dCTP, dGTP, and dTTP) at 37°C.

Terminal transferase assays were carried out in reaction medium containing 100 nM PPRT and 20 nM 5’-^32^P-labeled 20-nt single-stranded DNA oligonucleotide (3’-TG, Table S4) with 1 mM dNTPs at 37℃ for times up to 60 min.

Template-independent polymerization assays were done in reaction medium containing 200 nM PPRT with 50 μM dNTPs and 83 nM [α-^32^P]-dCTP at 37℃ for times up to 30 min.

Protein-priming assays were carried out in 20-μl reaction medium containing 500 nM PPRT at 37°C for 30 sec. After addition of dNTPs at a final concentration of 50 μM, reactions were incubated at 37°C for 15 min. Reactions were quenched with 4 μl of either (i) 125 mM EDTA, (ii) 125 mM EDTA and 200 U/ml Proteinase K, or (iii) 500,000 U/ml Micrococcal nuclease (MNase, New England Biolabs) and incubated for 30 min at 37°C. 4x NuPAGE Sample Buffer (Invitrogen) was added to a final 1x concentration and heated to 95°C for 3 min prior to electrophoresis. Samples were run on NuPAGE 4-12 % Bis-Tris gel, which was stained with Coomassie blue and scanned with a phosphorimager (Typhoon FL A 9500, GE Healthcare).

Misincorporation assays were carried out in reaction medium containing 100 nM PPRT, 22 nM 3’-Inv(dT)-blocked 50-nt DNA template (5’-TC or 5’-AC; Table S4), and 20 nM 5’-^32^P-labeled 20-nt DNA primer ending with a 3’-AG (Table S4), fully complementary to the 5’-TC template and with a 3’ mismatch for the 5’-AC template, with increasing concentration of dTTP or dATP (0.001, 0.01, 0.1, 1 mM) at 37℃ for 30 min.

Exonuclease assays were carried out in reaction medium containing 100 nM PPRT and 20 nM 5’-^32^P-labeled 20-nt DNA primer (3’-AG, Table S5) without or annealed to a 22 nM 3’-Inv(dT)-blocked 50-nt DNA template (Table S4) at 37℃ for times up to 120 min.

Translesion assays were carried out in reaction medium containing 100 nM PPRT, 22 nM 3’-Inv(dT)-blocked 50-nt template DNAs (Table S4) having a dG residue, 8-oxoguanine, or an AP site at position 23 nt from the 3’ end, and 20 nM 5’-^32^P-labeled 20-nt DNA primer (3’-TG, Table S4) with 1 mM dNTPs at 37℃ for times up to 60 min.

8-oxo-dG fidelity assays were carried out in reaction medium containing 100 nM PPRT, 22 nM 3’-Inv(dT)-blocked 50-nt template DNAs (Table S4) with an unmodified dG residue or an 8-oxo-dG lesion positioned 20 nt from the 3’-end, and 20 nM 5’-^32^P-labeled 19-nt DNA primer (3’-G, Table S4) at 37℃ for times up to 60 min.

To characterize phage defense DNAs synthesized by the RT-like domain, the PPRT protein and DNA products were treated with varying combinations of RNase A/T1 (ThermoFisher), thermolabile Proteinase K, nuclease P1, Duplex DNase, and MNase (all obtained from New England Biolabs). PPRT protein samples were prepared as control samples (-R; 1-μl storage buffer + 25 μl of 1 μM PPRT) or RNase A/T1 pre-incubated samples (+R; 1-μl 2500 U/ml RNase A/T1 + 25 μl of 1 μM PPRT) without subsequent RNase inactivation. Subsequent polymerization assays in the absence of added template were carried out in 20-μl reaction medium containing 4-μl -R or +R sample, 50 μM dNTPs, and 83 nM [α-^32^P]-dCTP at 37°C for 30 min. Reactions were quenched with 2 μl of 0.5 M EDTA or 120 U/ml thermolabile Proteinase K, followed by incubation at 37°C for 30 min. All samples were then incubated at 55°C for 10 min to inactivate the thermolabile Proteinase K, followed by 5 min room temperature equilibration. Samples were subsequently treated with 4 μl of either (i) 500,000 U/ml nuclease P1 in 5x NEBuffer 1.1, (ii) 1,000 U/ml Duplex DNase in 5x NEBuffer r2.1, or (iii) 1,000,000 U/ml MNase in 5x Micrococcal Nuclease Reaction Buffer (New England Biolabs) and incubated at 37°C for 20 min. After adding 13 μl of 3x formamide loading dye, the mixture was denatured by heating to 95°C for 3 min prior and DNA products were analyzed in an 8 M urea/6% polyacrylamide gel run at 55 W for 1.5 h. The gel was then dried and scanned with a phosphorimager (Typhoon FL A 9500, GE Healthcare).

Thin layer chromatography (TLC) assays were carried out with in reaction medium containing 100 nM PPRT and 20 nM 3’-^32^P-labeled 20-nt DNA primer (3’-TG, Table S4) with 1 mM dNTPs at 37°C for 30 min. After termination of reactions using STOP solution, the samples were run on PEI (polyethyleneimine)-cellulose thin layer chromatography (TLC) plate (Sigma-Aldrich) against untreated [α-^32^P]-dCTP and apyrase-treated [α-^32^P]-dCTP, which generates [α-^32^P]-dCDP and [α-^32^P]-dCMP, serving as markers in parallel lanes. in TLC running buffer (0.1 M potassium phosphate (KPi) pH 3.8) for 2 h. Plates were dried in room temperature and scanned with phosphorimager (Typhoon FL A 9500, GE Healthcare).

### Sequencing of DNA synthesized by PPRT without an added template

DNA synthesis reactions without an added template were done in 100-μl of reaction medium containing 200 nM enzyme with 50 µM dNTPs at 37°C for 30 min. 20-μl STOP solution was added to stop the reaction and incubated at 37°C for 30 min for protein digestion and release of protein-primed DNA products. DNA sequencing libraries were generated by TGIRT-seq protocol and sequenced on a MiSeq v3 instrument to obtain ~1 million 2 x 300 nt paired end reads per sample at the GSAF at the University of Texas at Austin. Illumina TruSeq adapters and PCR primer sequences were trimmed from the reads with Cutadapt v4.9 (https://github.com/marcelm/cutadapt, sequencing quality score cut-off at 20; p-value <0.01), and reads <15 nt after trimming were discarded. Trimmed reads were filtered by Kraken v2.1.3^81^ using the Standard PlusPF database (https://benlangmead.github.io/aws-indexes/k2)^81^ to remove potential contaminant reads derived from Human, RefSeq archaea, bacteria, protozoa, fungi, viral, plasmid, and UniVec_core sequences. Paired-end reads were then merged by fastp v0.23.4 (https://github.com/OpenGene/fastp)^82^ using the following parameters “-A -m -c -overlap_len require 10”. Merged reads were then checked by FastQC v0.12.1 (https://github.com/s-andrews/FastQC)^83^ to identify overrepresented sequences. One short sequence “TGGGAAGCTCAGAATAAACGCTCAACTTTGG” was detected in all 4 datasets. A Blast search of this sequence found that it matches to part of the 5’ UTR used in many protein expression vectors. Additional steps were taken to remove this overrepresented sequence by mapping merged reads against it using Hisat2 v2.2.1 (http://daehwankimlab.github.io/hisat2)^84^. Unmapped reads corresponding to DNAs synthesized by PPRT without an added template were collected and analyzed to determine nucleotide frequencies and the lengths of homopolymer runs by using R v4.3.3 (https://www.r-project.org)^77^.

### Primer extension assays and DNA sequencing of products with matched and mismatched 3’-primer nucleotides

Primer extension assays comparing template-primer combinations with matched and mismatched 3’-primer nucleotides were carried out as above in 100-μl reaction medium containing 100 nM PPRT, 22 nM 3’-Inv(dT)-blocked 50-nt DNA (5’-AC or 5’-TC, Table S4), and 20 nM 20-nt DNA primer (3’-AG, Table S4) at 37°C for 2 h. Reactions were terminated by adding 20-μl STOP solution (see above) and incubated at 37°C for 30 min for proteinase K digestion to release of protein-primed DNA products. DNA products were used for generating DNA sequencing libraries using the TGIRT-seq protocol and sequenced on the MiSeq v3 instrument to obtain ~1 million 2 x 300 nt paired end reads per sample at the GSAF at the University of Texas at Austin. Illumina TruSeq adapters and PCR primer sequences were trimmed from the reads with Cutadapt v4.9 (sequencing quality score cut-off at 20; p-value <0.01) and reads <15 nt after trimming were discarded. Trimmed reads were merged by fastp using the same parameters indicated above. Merged reads were then mapped to the DNA template sequence and mapped reads were retrieved and realigned to the template sequence by MAFFT v7.526 (https://github.com/GSLBiotech/mafft)^85^.

