## Supplementary File for "A bacterial PrimPol-reverse transcriptase hybrid protein has a proofreading exonuclease activity that can be transferred to other reverse transcriptases"

This PDF file includes:
Figures S1 to S5
Tables S1 to S6

**Figure S1**

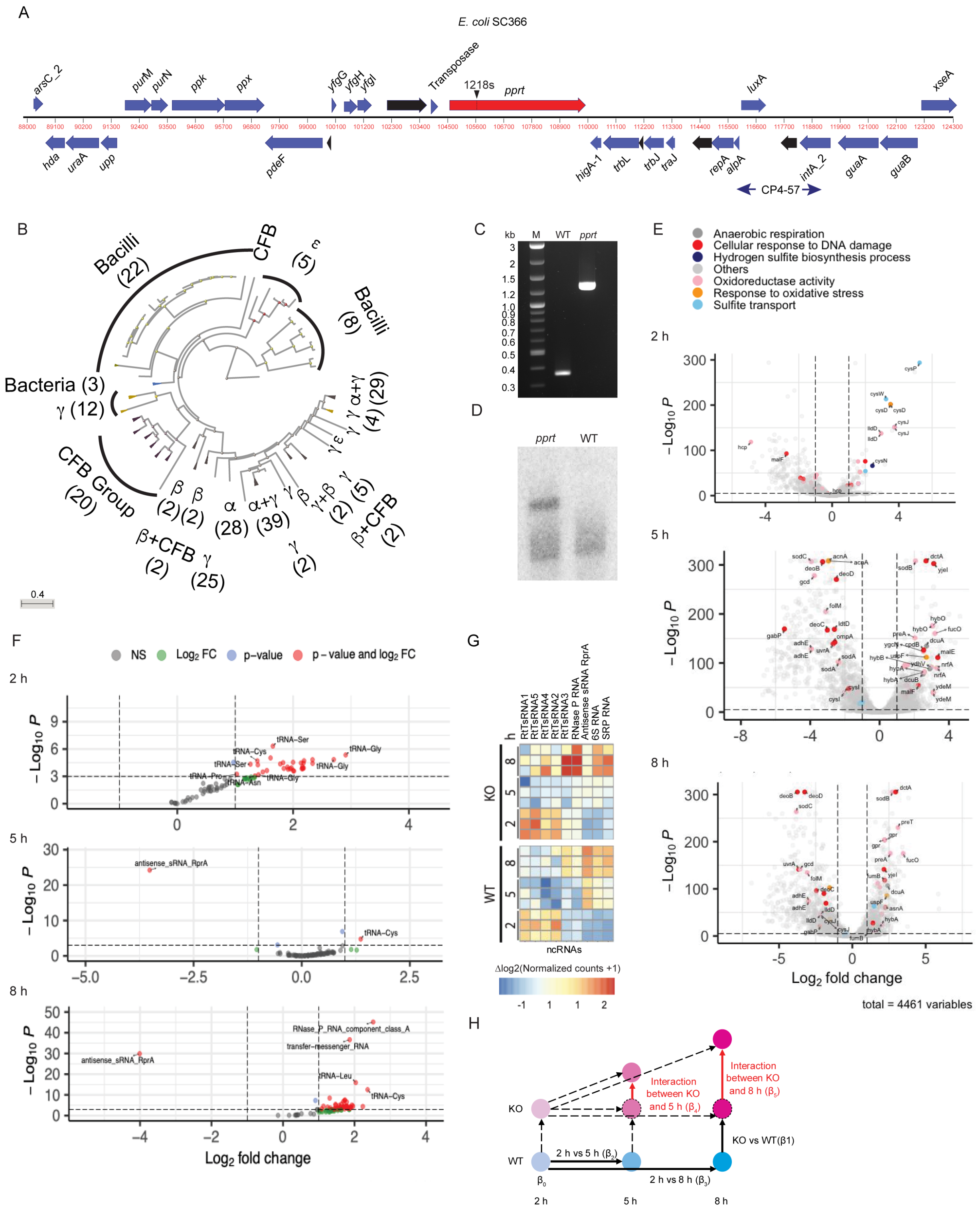

A
E. coli SC366
arsC_2
purM
purN
ppk
ppx
yfgG
yfgH
yfgI
Transposase
1218s
pprt
luxA
xseA
hda
uraA
upp
pdeF
higA-1
trbL
trbJ
traJ
repA
alpA
intA_2
guaA
guaB
CP4-57
B
Bacilli (22)
CFB
Bacilli (8)
Bacteria (3)
γ (12)
CFB Group (20)
β+CFB (2)
γ (25)
α (28)
α+γ (39)
β+CFB (2)
0.4
C
kb
M
WT
pprt
D
pprt
WT
E
Anaerobic respiration
Cellular response to DNA damage
Hydrogen sulfite biosynthesis process
Others
Oxidoreductase activity
Response to oxidative stress
Sulfite transport
2 h
5 h
8 h
Log2 fold change
total = 4461 variables
F
NS
Log2 FC
p−value
p − value and log2 FC
2 h
5 h
8 h
antisense_sRNA_RprA
RNase_P_RNA_component_class_A
transfer−messenger_RNA
tRNA−Leu
tRNA−Cys
tRNA−Ser
tRNA−Gly
Log2 fold change
G
RtTsRNA1
RtTsRNA5
RtTsRNA4
RtTsRNA2
RtTsRNA3
RNase P RNA
Antisense sRNA RprA
6S RNA
SRP RNA
KO
WT
ncRNAs
Δlog2(Normalized counts +1)
H
KO
WT
Interaction between KO and 5 h (β4)
Interaction between KO and 8 h (β5)
KO vs WT(β1)
2 h vs 5 h (β2)
2 h vs 8 h (β3)
β0
2 h
5 h
8 h

**Figure S1. *E. coli* SC366 PPRT gene region and characteristics**

(A) Map of the *E. coli* SC366 genome region encoding PPRT (contig QOMS01000055). The *pprt* gene is shown as a red arrow with the targetron insertion site indicated by an arrowhead at 1218s (sense-strand nucleotide 1218 of *pprt*). Genes encoding proteins with annotated functions are shown as blue arrows, and genes encoding proteins without known functions are shown as black arrows. Red numbers indicate nucleotide positions in the *E. coli* SC366 genome (QOMS01000055). A CP4-57 prophage-derived region is indicated below by a double-headed arrow.

(B) Phylogram based on ~200 *pprt* gene sequences identified by a BLASTP search using the *E. coli* SC366 *pprt* gene sequence as a query. The phylogram was generated by the BLAST tree viewer using default settings. $\alpha$, $\beta$, $\gamma$, and $\varepsilon$ refer to different groups of proteobacteria, with the numbers in parentheses indicating the number of sequences for different groups. CFB, gram negative Cytophaga-Flavobacterium-Bacteroides group.

(C) Verification of targetron insertion at the desired site by PCR of extracted genomic DNA from the *pprt* disruptant with primers 1218-F and 1218-R with the product run on a 1.0% agarose gel against a 1 kb plus DNA ladder (New England Biolabs).

(D) Southern hybridization showing targetron insertion in the *pprt* gene. Genomic DNAs from WT and *pprt* disruptant strains were digested with HindIII and EcoRI, separated on 1.0% agarose gel, and transferred to nylon membrane. Blots were hybridized with a $^{32}$P-labeled 197-bp targetron probe (Table S6).

(E) Pairwise volcano plots comparing differentially expressed protein-coding genes in the *pprt* disruptant versus WT strain at 2, 5, and 8 h. Horizontal dashed lines indicate the p-value significance threshold, while vertical dashed lines mark the significant fold-change cutoffs.

(F) Volcano plots comparing differentially expressed non-coding RNAs in the *pprt* disruptant versus WT strain at 2, 5, and 8 h. Gray circles, non-significantly differentially expressed genes; green circles, genes with significant $\log_2$ fold change only; blue circles, genes with significant p-values only; red circles, genes meeting both p-value and $\log_2$ fold change significance criteria. Horizontal dashed lines indicate the p-value significance threshold, while vertical dashed lines mark the fold change cutoffs.

(G) Heat map comparing differentially expressed sncRNAs in the *pprt* disruptant versus WT strain at 2, 5, and 8 h.

(H) Model for designing the matrix used in DESeq2 analyses. The model considers the effect of differences in genotype, growth times, and the interactions between genotype and growth time differences. The blue spheres represent the WT genotype, and the purple spheres represent the *pprt* KO genotype. The light-dark contrast reflects the time points, with darker contrast emphasizing the genotype specific effects accumulated in the cells at indicated time points. The sphere with the dashed outline is an imaginary condition for the linear addition of the difference due to the genotype and growth, which is used as the background to understand the interactions between the genotype and growth time differences. Solid black arrows indicate direct comparisons corresponding to model coefficients: $\beta_0$ (baseline, wild type 2 h), $\beta_1$ (genotype effect, KO vs WT), $\beta_2$ (time differences, 2 h vs 5 h in WT), and $\beta_3$ (time differences, 2 h vs 8 h in WT). Dashed black lines represent expected additive effects under linear model assumptions, showing theoretical KO positions without interaction effects. Red arrows indicate interaction terms $\beta_4$ (KO×5 h interaction) and $\beta_5$ (KO×8 h interaction), measuring deviations from expected additive effects to capture genotype-time dependencies.

### Figure S2

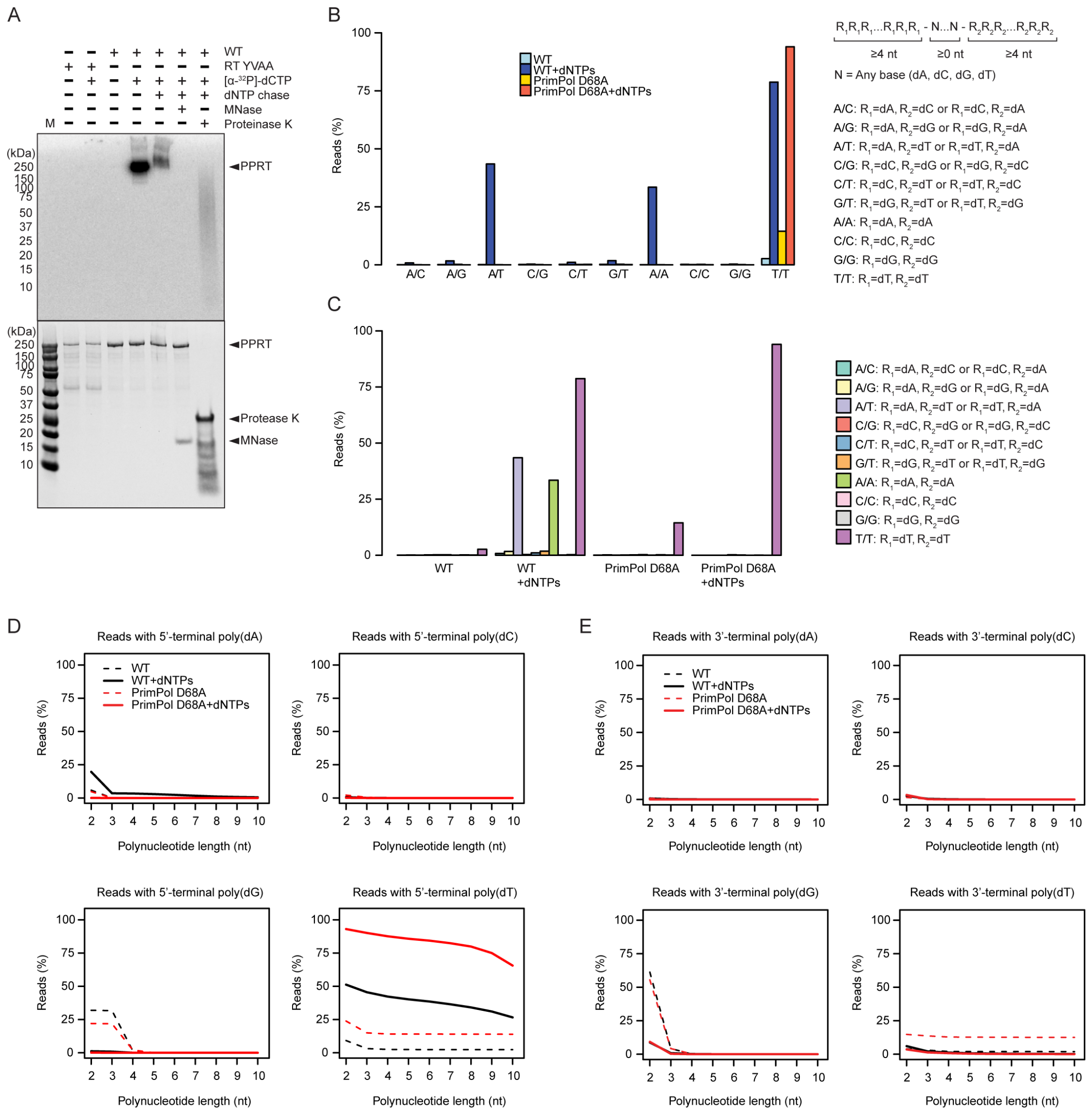
A
WT
RT YVAA
[α-32P]-dCTP
dNTP chase
MNase
Proteinase K
M
(kDa)
250
150
100
75
50
37
25
20
15
10
◄PPRT
◄Protease K
◄MNase
B
WT
WT+dNTPs
PrimPol D68A
PrimPol D68A+dNTPs
Reads (%)
A/C
A/G
A/T
C/G
C/T
G/T
A/A
C/C
G/G
T/T
R1R1R1...R1R1R1 - N...N - R2R2R2...R2R2R2
≥4 nt
≥0 nt
≥4 nt
N = Any base (dA, dC, dG, dT)
A/C: R1=dA, R2=dC or R1=dC, R2=dA
A/G: R1=dA, R2=dG or R1=dG, R2=dA
A/T: R1=dA, R2=dT or R1=dT, R2=dA
C/G: R1=dC, R2=dG or R1=dG, R2=dC
C/T: R1=dC, R2=dT or R1=dT, R2=dC
G/T: R1=dG, R2=dT or R1=dT, R2=dG
A/A: R1=dA, R2=dA
C/C: R1=dC, R2=dC
G/G: R1=dG, R2=dG
T/T: R1=dT, R2=dT
C
Reads (%)
WT
WT +dNTPs
PrimPol D68A
PrimPol D68A +dNTPs
D
Reads with 5'-terminal poly(dA)
Reads with 5'-terminal poly(dC)
Reads with 5'-terminal poly(dG)
Reads with 5'-terminal poly(dT)
WT
WT+dNTPs
PrimPol D68A
PrimPol D68A+dNTPs
Reads (%)
Polynucleotide length (nt)
E
Reads with 3'-terminal poly(dA)
Reads with 3'-terminal poly(dC)
Reads with 3'-terminal poly(dG)
Reads with 3'-terminal poly(dT)

**Figure S2. PPRT protein-primed DNA synthesis**

(A) Protein-priming assays for WT and RT YVAA with 50 µM dNTPs plus 83 nM [α-$^{32}$P]-dCTP in reaction medium containing 500 nM enzyme, 10 mM NaCl, 10 mM $MgCl_2$, 1 mM $MnCl_2$, and 20 mM Tris-HCl pH 7.5 at 37°C for 30 sec, followed by incubation of samples with (i) 125 mM EDTA, (ii) 125 mM EDTA and 200 U/ml Proteinase K, or (iii) 500,000 U/ml Micrococcal Nuclease (MNase) and further incubation at 37°C for 30 min. Samples were run on NuPAGE 4-12% Bis-Tris gel and stained with Coomassie blue. The gel was scanned with a phosphorimager (Typhoon FL A 9500, GE Healthcare). This analysis showed that [α-$^{32}$P]-dCTP forms a covalent bond, resulting in labeled protein in an SDS-polyacrylamide gel and that a chase with unlabeled dNTPs shifts the labeled WT protein band upward in the gel, indicating that the DNA was elongated while covalently attached to PPRT. MNase treatment completely removed all traces of radiolabeled DNA from PPRT, as expected for DNA products, while Proteinase K treatment resulted in a smear of labeled bands that ranged from 10 to 250 kDa, as expected for DNA products of different length bound to protein fragments. The PPRT YVAA mutant showed no labeled bands, confirming that the RT active site accounts for the protein-priming activity.

(B and C) Distribution of patterns of single nucleotide runs of different combinations in ssDNAs synthesized by WT and D68A PPRT proteins. Template-independent DNA polymerization assays were carried out with 100 nM WT and mutant PPRTs and 50 µM dNTPs in the absence of DNA templates or primers in reaction medium containing 100 nM enzyme, 10 mM NaCl, 10 mM $MgCl_2$, 1 mM $MnCl_2$, and 20 mM Tris-HCl pH 7.5 at 37°C for times up to 30 min (see Methods). Illumina MiSeq sequencing showed the percentage of reads having runs of the same nucleotide ($R1 \geq 4$ nt) followed by runs of the same or different nucleotides ($R2 \geq 4$ nt) for WT PPRT without or with added dNTPs (light and dark blue bars, respectively) and PrimPol D68A without or with added dNTPs (yellow and red bars, respectively), organized by combinations of Poly(dN) patterns (B) or experimental conditions (C), respectively. Combinations of Poly(dN) patterns shown to the right indicate the specific homopolymer and repetitive sequences with their corresponding sequence contexts; $R_1$, N, and $R_2$.

(D and E) Plots showing the MiSeq read length distribution of the synthesized ssDNA products (nt) by WT and PrimPol D68A mutant that start (5'-terminal) or end (3'-terminal) with poly(dA), poly(dG), poly(dC), or poly(dT).

**Figure S3**

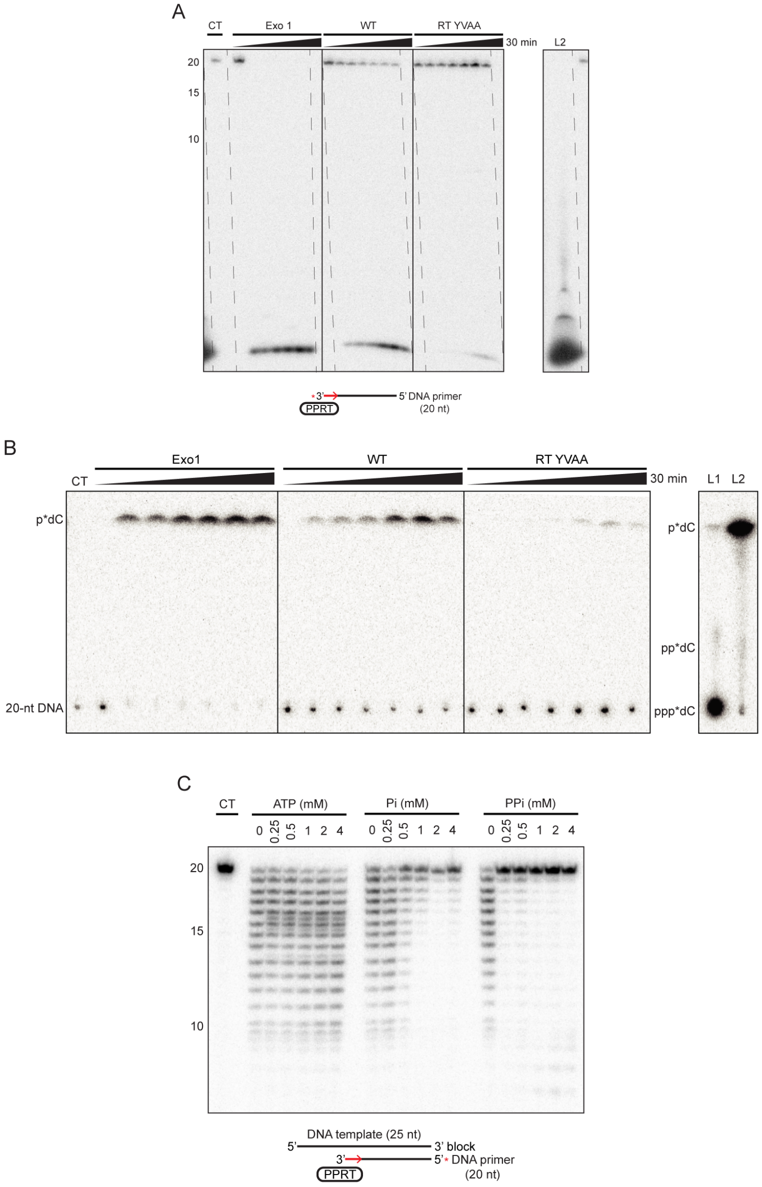
A
CT
Exo 1
WT
RT YVAA
30 min
L2
20
15
10
3'
5' DNA primer
(20 nt)
PPRT
B
Exo1
WT
RT YVAA
30 min
CT
L1
L2
p*dC
p*dC
pp*dC
20-nt DNA
ppp*dC
C
CT
ATP (mM)
Pi (mM)
PPi (mM)
0 0.25 0.5 1 2 4
0 0.25 0.5 1 2 4
0 0.25 0.5 1 2 4
20
15
10
DNA template (25 nt)
5'
3' block
3'
5' DNA primer
(20 nt)
PPRT

**Figure S3. PPRT exonuclease activity**

(A and B) Exonuclease I (Exo I, New England Biolabs), WT PPRT, and YVAA mutant PPRT incubated with a 3'-$^{32}$P-labeled (red asterisk) 20-nt DNA primer in reaction medium containing 100 nM enzyme, 10 mM NaCl, 10 mM $MgCl_2$, 1 mM $MnCl_2$, and 20 mM Tris-HCl pH 7.5 at 37°C for times up to 15 min and run either on a denaturing 17% polyacrylamide gel against 5'-labeled oligonucleotide size markers in a parallel lane (see Methods) (panel A) or a polyethyleneimine-cellulose thin layer chromatography plate against untreated [α-$^{32}$P]-dCTP (L1; Ladder 1) and apyrase-treated [α-$^{32}$P]-dCTP (L2; Ladder 2), which generates [α-$^{32}$P]-dCDP and [α-$^{32}$P]-dCMP, serving as markers in parallel lanes (panel B). CT (control) was tested without enzyme under identical conditions, with an incubation time of 30 min.

(C) Exonuclease assay of WT PPRT using a 3'-Inv(dT)-blocked 25-nt DNA template and a 3'-$^{32}$P-labeled (red asterisk) 20-nt DNA primer in the presence of different concentrations of ATP, Pi, and PPi. The reactions were done in the same reaction medium as panels A and B at 37°C for times up to 60 min. CT (control) was tested without enzyme and inorganic phosphates under identical conditions, with an incubation time of 60 min. The DNA products were analyzed in a 17% polyacrylamide gel against 5'-labeled oligonucleotide size markers in a parallel lane (see Methods).

**Figure S4**

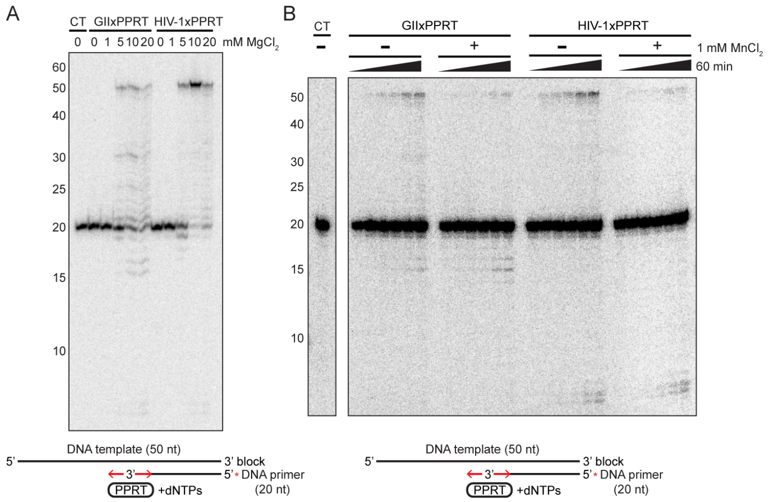

**Figure S4. $Mg^{2+}$- and $Mn^{2+}$-dependence of primer extension and exonuclease activities of hybrid GIIxPPRT and HIV-1xPPRT.**

(A) Primer extension assays of GIIxPPRT and HIV-1xPPRT assayed using a 3'-Inv(dT)-blocked 50-nt DNA template and a 5'-$^{32}$P-labeled (red asterisk) 20-nt DNA primer done in reaction medium containing 100 nM enzyme, 10 mM NaCl, different concentrations of $MgCl_2$, and 20 mM Tris-HCl pH 7.5 at 37°C for 60 min. DNA products were analyzed in a 17% polyacrylamide gel against 5'-labeled oligonucleotide size markers in a parallel lane (see Methods). CT (control) was tested without enzyme under identical conditions, with an incubation time of 60 min.

(B) Primer extension assays of GIIxPPRT and HIV-1xPPRT were done as in panel A in reaction medium containing 100 nM enzyme, 10 mM NaCl, 10 mM $MgCl_2$, without (-) or with (+) 1 mM $MnCl_2$, and 20 mM Tris-HCl pH 7.5 at 37°C for times up to 60 min. CT (control) was tested without enzyme under identical conditions, with an incubation time of 60 min.

**Figure S5**

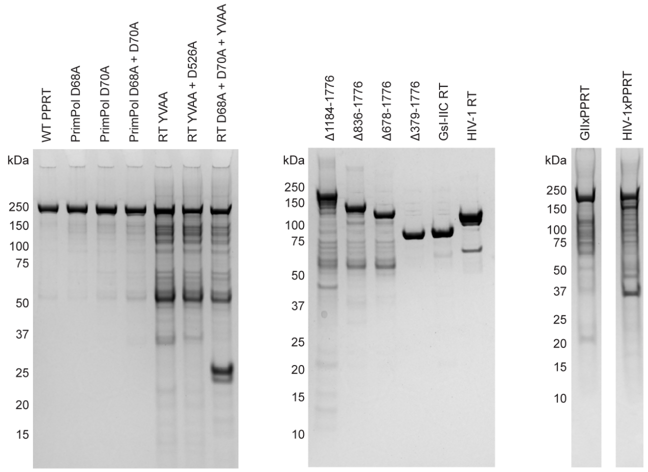

**Figure S5. Purified PPRT proteins analyzed by SDS-polyacrylamide electrophoresis.** Protein samples (0.5 µg**)** were analyzed by electrophoresis in a NuPAGE 4-12% Bis-Tris gel and stained with Coomassie blue against prestained protein standard (10-250 kDa; New England Biolabs).

**Table S1. TGIRT-seq detected mutation patterns in WT and *pprt* disruptant strain.**

| Wild Type | Disruptant | WT_2 h | WT_5 h | WT_8 h | KO_2 h | KO_5 h | KO_8 h | |
|---|---|---|---|---|---|---|---|---|
| | | 33.33 | 31.67 | 50.33 | 44.33 | 36.00 | 62.00 | AVG |
| A | C | 4.62 | 4.04 | 8.02 | 3.79 | 2.00 | 17.44 | STDEV |
| | | 5.23 | 4.57 | 9.08 | 4.28 | 2.26 | 19.73 | Confidence Interval (95%) |
| | | | | | 0.03 | 0.17 | 0.35 | t-test (time-wise) |
| | | 144.33 | 148.67 | 163.00 | 144.33 | 173.67 | 164.00 | AVG |
| A | G | 19.50 | 14.50 | 23.81 | 3.51 | 15.57 | 22.34 | STDEV |
| | | 22.07 | 16.41 | 26.95 | 3.97 | 17.62 | 25.28 | Confidence Interval (95%) |
| | | | | | 1.00 | 0.11 | 0.96 | t-test (time-wise) |
| | | 41.67 | 36.00 | 53.00 | 48.67 | 49.67 | 54.67 | AVG |
| A | T | 6.03 | 1.00 | 3.00 | 10.60 | 8.50 | 7.51 | STDEV |
| | | 6.82 | 1.13 | 3.39 | 11.99 | 9.62 | 8.49 | Confidence Interval (95%) |
| | | | | | 0.38 | 0.05 | 0.74 | t-test (time-wise) |
| C | A | 191.00 | 165.00 | 232.00 | 191.33 | 180.33 | 141.67 | AVG |
| | | 70.87 | 10.15 | 64.63 | 6.11 | 11.02 | 17.62 | STDEV |
| | | 80.20 | 11.48 | 73.13 | 6.91 | 12.46 | 19.93 | Confidence Interval (95%) |
| | | | | | 0.99 | 0.15 | 0.08 | t-test (time-wise) |
| | | 46.33 | 47.33 | 57.67 | 64.00 | 61.00 | 53.33 | AVG |
| C | G | 1.53 | 6.51 | 3.79 | 11.53 | 4.00 | 2.08 | STDEV |
| | | 1.73 | 7.36 | 4.28 | 13.05 | 4.53 | 2.36 | Confidence Interval (95%) |
| | | | | | 0.06 | 0.04 | 0.16 | t-test (time-wise) |
| | | 218.67 | 199.67 | 221.33 | 204.67 | 227.33 | 188.33 | AVG |

| | | | | | | | | |
|---|---|---|---|---|---|---|---|---|
| C | T | 21.94 | 40.41 | 17.04 | 4.04 | 21.57 | 15.57 | STDEV |
| | | 24.83 | 45.73 | 19.28 | 4.57 | 24.41 | 17.62 | Confidence Interval (95%) |
| | | | | | 0.34 | 0.35 | 0.07 | t-test (time-wise) |
| | | 219.33 | 192.33 | 232.67 | 209.67 | 237.67 | 188.00 | AVG |
| G | A | 12.58 | 17.16 | 12.01 | 9.61 | 1.53 | 29.87 | STDEV |
| | | 14.24 | 19.41 | 13.59 | 10.87 | 1.73 | 33.80 | Confidence Interval (95%) |
| | | | | | 0.35 | 0.01 | 0.07 | t-test (time-wise) |
| | | 47.67 | 43.00 | 64.33 | 61.33 | 50.67 | 55.67 | AVG |
| G | C | 5.69 | 2.65 | 3.51 | 10.41 | 3.51 | 20.11 | STDEV |
| | | 6.43 | 2.99 | 3.97 | 11.78 | 3.97 | 22.75 | Confidence Interval (95%) |
| | | | | | 0.12 | 0.04 | 0.50 | t-test (time-wise) |
| | | 199.00 | 165.67 | 245.33 | 174.67 | 180.00 | 146.00 | AVG |
| G | T | 60.56 | 16.04 | 90.69 | 17.01 | 32.08 | 13.86 | STDEV |
| | | 68.52 | 18.15 | 102.63 | 19.25 | 36.30 | 15.68 | Confidence Interval (95%) |
| | | | | | 0.54 | 0.53 | 0.13 | t-test (time-wise) |
| | | 39.67 | 41.33 | 43.33 | 48.00 | 52.67 | 41.33 | AVG |
| T | A | 5.86 | 5.51 | 3.06 | 5.29 | 3.06 | 1.53 | STDEV |
| | | 6.63 | 6.23 | 3.46 | 5.99 | 3.46 | 1.73 | Confidence Interval (95%) |
| | | | | | 0.14 | 0.04 | 0.37 | t-test (time-wise) |
| | | 137.67 | 145.00 | 168.00 | 133.00 | 164.33 | 167.33 | AVG |
| T | C | 17.16 | 9.54 | 13.86 | 3.61 | 27.54 | 13.01 | STDEV |
| | | 19.41 | 10.79 | 15.68 | 4.08 | 31.16 | 14.73 | Confidence Interval (95%) |
| | | | | | 0.67 | 0.31 | 0.95 | t-test (time-wise) |

| | | | | | | | | |
|---|---|---|---|---|---|---|---|---|
| | | 39.33 | 35.00 | 46.33 | 46.00 | 44.67 | 48.33 | AVG |
| T | G | 13.01 | 3.00 | 12.58 | 5.57 | 2.52 | 6.35 | STDEV |
| | | 14.73 | 3.39 | 14.24 | 6.30 | 2.85 | 7.19 | Confidence Interval (95%) |
| | | | | | 0.46 | 0.01 | 0.82 | t-test (time-wise) |

In the WT strain, mutation levels decreased at 5 h followed by an increase at 8 h. Conversely, in the *pprt* disruptant strain mutation levels increased at 5 h followed by a decrease at 8 h prior to early entry into stationary phase (Figure 1K). The predominant mutation types observed were A to T, T to A, G to A, and T to G mutations. p-values ≤0.05 indicated in red.

**Table S2. List of *E. coli* strains used.**

| Strain | Source | Source or reference |
|---|---|---|
| DH5α | Clone strain, F– φ80lacZΔM15 Δ(lacZYA-argF)U169 recA1 endA1 hsdR17(rK–, mK+) phoA supE44 λ–thi-1 gyrA96 relA1 | ThermoFisher |
| Rosetta2(DE3) | Expression strain, F- ompT hsdSB(rB- mB-) gal dcm (DE3) pRARE2 (CamR) | Novagen |
| Rosetta2(DE3)::pMal-GII RT | Derived from Rosetta2(DE3), carrying expression plasmid pMal-GII RT | This study |
| Rosetta2(DE3)::pMal-GIIxPPRT | Derived from Rosetta2(DE3), carrying expression plasmid pMal-GIIxPPRT | This study |
| Rosetta2(DE3)::pMal-HIV-1 RT | Derived from Rosetta2(DE3), carrying expression plasmid pMal-HIV-1 RT | This study |
| Rosetta2(DE3)::pMal-HIV-1xPPRT | Derived from Rosetta2(DE3), carrying expression plasmid pMal-HIV-1xPPRT | This study |
| Rosetta2(DE3)::pMal-PPRT | Derived from Rosetta2(DE3), carrying expression plasmid pMal-PRT | This study |
| Rosetta2(DE3)::pMal-PPRT-D526A+YVAA | Derived from Rosetta2(DE3), carrying expression plasmid pMal-PPRT-D526A+YVAA | This study |
| Rosetta2(DE3)::pMal-PPRT-D68A | Derived from Rosetta2(DE3), carrying expression plasmid pMal-PPRT-D68A | This study |
| Rosetta2(DE3)::pMal-PPRT-D68A+D70A | Derived from Rosetta2(DE3), carrying expression plasmid pMal-PPRT-D68A+D70A | This study |
| Rosetta2(DE3)::pMal-PPRT-D68A+D70A+YVAA | Derived from Rosetta2(DE3), carrying expression plasmid pMal-PPRT-D68A+D70A+YVAA | This study |
| Rosetta2(DE3)::pMal-PPRT-D70A | Derived from Rosetta2(DE3), carrying expression plasmid pMal-PPRT-D70A | This study |
| Rosetta2(DE3)::pMal-PPRT-YVAA | Derived from Rosetta2(DE3), carrying expression plasmid pMal-PPRT-YVAA | This study |
| Rosetta2(DE3)::pMal-PPRTΔ1184-1776 | Derived from Rosetta2(DE3), carrying expression plasmid pMal-PPRTD1184-1776 | This study |
| Rosetta2(DE3)::pMal-PPRTΔ379-1776 | Derived from Rosetta2(DE3), carrying expression plasmid pMal-PPRTD379-1776 | This study |
| Rosetta2(DE3)::pMal-PPRTΔ678-1776 | Derived from Rosetta2(DE3), carrying expression plasmid pMal-PPRTD678-1776 | This study |
| Rosetta2(DE3)::pMal-PPRTΔ836-1776 | Derived from Rosetta2(DE3), carrying expression plasmid pMal-PPRTD836-1776 | This study |

| | | |
|---|---|---|
| Top 10 | F⁻mcrA Δ(mrr-hsdRMS-mcrBC) φ80lacZΔM15 ΔlacX74 recA1 araD139 Δ(ara-leu)7697 galU galK λ⁻rpsL($Str^R$) endA1 nupG | ThermoFisher |

**Table S3. List of plasmids used.**

| **Plasmid** | **Source** |
|---|---|
| pBL1 | Yao, J., & Lambowitz, A. M. (2007). Applied and environmental microbiology, 73(8), 2735-2743. |
| pBL1-PPRT::1218s | This study |
| pCR$^{TM}$2.1 T vector | ThermoFisher |
| pMal-c5X | New England Biolabs |
| pMal-GII RT | Mohr, S., Ghanem, E., Smith, W., Sheeter, D., Qin, Y., King, O., ... & Lambowitz, A. M. (2013). RNA, *19*(7), 958-970. |
| pMal-GIIxPPRT | This study |
| pMal-HIV-1 RT | This study |
| pMal-HIV-1xPPRT | GenScript |
| pMal-PPRT | This study |
| pMal-PPRT-D526A+YVAA | This study |
| pMal-PPRT-D68A | This study |
| pMal-PPRT-D68A+D70A | This study |
| pMal-PPRT-D68A+D70A+YVAA | This study |
| pMal-PPRT-D70A | This study |
| pMal-PPRT-YVAA | This study |
| pMal-PPRTΔ1184-1776 | This study |
| pMal-PPRTΔ379-1776 | This study |
| pMal-PPRTΔ678-1776 | This study |
| pMal-PPRTΔ836-1776 | This study |

**Table S4. List of DNA and RNA substrates used.**

| **Oligonucleotides** | **Sequence (5' to 3')** |
|---|---|
| 19-nt DNA primer (3'-G) | TTGGGACCTTAAGAATTTG |
| 20-nt DNA primer (3'-AG) | TTGGGACCTTAAGAATTTGA |
| 20-nt DNA primer (3'-TG) | TTGGGACCTTAAGAATTTGT |
| 25-nt DNA 3' block (5'-AC) | TACAAACAAATTCTTAAGGTCCCAA/Inv(dT)/ |
| 50-nt DNA 3' block (5'-AC) | GCAATAATCTATACAATACAACACATACAAACAAATTCT TAAGGTCCCAA/Inv(dT)/ |
| 50-nt DNA 3' block (5'-AG) | GCAATAATCTATACAATACAACACATACAAAGAAATTCT TAAGGTCCCAA/Inv(dT)/ |
| 50-nt DNA 3' block (5'-TC) | GCAATAATCTATACAATACAACACATACAATCAAATTCT TAAGGTCCCAA/Inv(dT)/ |
| 50-nt RNA 3' block | GCAAUAAUCUAUACAAUACAACACAUACAAACAAAUUC UUAAGGUCCCAA/Inv(dT)/ |
| 8-oxoguanine 50-nt DNA 3' block (8-oxo-dG positioned 20 nt from 3' end) | GCAATAATCTATACAATACAACACATACAA/8oxodG/CAA ATTCTTAAGGTCCCAA/Inv(dT)/ |
| 8-oxoguanine 50-nt DNA 3' block (8-oxo-dG positioned 23 nt from 3' end) | GCAATAATCTATACAATACAACACATA/8oxodG/AAACAA ATTCTTAAGGTCCCAA/Inv(dT)/ |
| AP site 50-nt DNA 3' block | GCAATAATCTATACAATACAACACATA/idSp/AAACAAAT TCTTAAGGTCCCAA/Inv(dT)/ |
| dG residue 50-nt DNA 3' block | GCAATAATCTATACAATACAACACATAGAAACAAATTCT TAAGGTCCCAA/Inv(dT)/ |

Major sequence differences are underlined. Abbreviations: Inv(dT), inverted dT; idSp, internal 1',2'-dideoxyribose spacer (dSpacer).

**Table S5. List of primers used.**

| Primer | Sequence (5' to 3') | Note |
|---|---|---|
| 0104-1 | CAACGGTACGTAGCAGCAATTTTCTTCGTAGCACCTAC | To construct D626A/D627A point mutation plasmid |
| 0104-2 | GAAAGAACATGTGAGCAAAAG | To construct D626A/D627A point mutation plasmid |
| 0104-3 | CTTTTGCTCACATGTTCTTTC | To construct D626A/D627A point mutation plasmid |
| 0104-4 | TATCATCTACGTACCGTTGG | To construct D626A/D627A point mutation plasmid |
| 1184-F | CTTATGAATTCGCTGAATGAGCATCATCATCATCATCATCACTAAC TGCAGGCAAG | To construct Δ1184-1776 disruption plasmid |
| 1184-R | CTTGCCTGCAGTTAGTGATGATGATGATGATGATGCTCATTCAGCG AATTCATAAG | To construct Δ1184-1776 disruption plasmid |
| 1218 EBS1d | CAGATTGTACAAATGTGGTGATAACAGATAAGTCAAAATCATTAA CTTACCTTTCTTTGT | To amplify the retargeted intron |
| 1218 EBS2 | TGAACGCAAGTTTCTAATTTCGGTTGACTGTCGATAGAGGAAAGTG TCT | To amplify the retargeted intron |
| 1218 EBSU | CGAAATTAGAAACTTGCGTTCAGTAAAC | To amplify the retargeted intron |
| 1218 IBS | AAAAAAGCTTATAATTATCCTTACAGTCCAAAATCGTGCGCCCAGA TAGGGTG | To amplify the retargeted intron |
| 1218-F | CCATCACAAGAGCAAATAG | To detect the intron insertion |
| 1218-R | AGCTATTCTCTCTGGTGC | To detect the intron insertion |
| 379-F | CATCATCATCATCATCATCACTAACTGCAG | To construct Δ379-1776 disruption plasmid |
| 379-R | ATCAAATGTAGCATCTGTATAAGCTAAAAATTTTGGTATGCA | To construct Δ379-1776 disruption plasmid |
| 678-F | CATCATCATCATCATCATCACTAACTGCAG | To construct Δ678-1776 disruption plasmid |
| 678-R | ACGATAACTGTTAAAATCTTCAGAGAAATTACTTATTTTTGTTATTA CT | To construct Δ678-1776 disruption plasmid |
| 836-F | CATCATCATCATCATCATCACTAACTGCAG | To construct Δ836-1776 disruption plasmid |

| | | |
|---|---|---|
| 836-R | GATTTTGGCATCGAACACCAATGC | To construct Δ836-1776 disruption plasmid |
| D526A_F | CGTAAAAACTGCGATCAGAAATTTTTATGACACG | To construct D526A point mutation plasmid |
| D526A_R | ATATAGCACTGTTGATTATCATTGTC | To construct D526A point mutation plasmid |
| D68A_2-F | GATCTGCTTTGCTTTTGACATAATTAAATCTAATTTAAATAG | To construct D68A+D70A+YVAA disruption plasmid |
| D68A_2-R | CAGTTAATTGTGAGATCGTTATTTTTTTG | To construct D68A+D70A+YVAA disruption plasmid |
| D68A-F | CACAATTAACTGGATCTGCTTTGCTTTTGACATAATTAAATCTAAT | To construct D68A point mutation plasmid |
| D68A-R | ATTAGATTTAATTATGTCAAAAGCAAAGCAGATCCAGTTAATTGTG | To construct D68A point mutation plasmid |
| D70A_2-F | CTTTGCTTTTGCTATAATTAAATCTAATTTAAATAG | To construct D68A+D70A and D68A+D70A+YVAA disruption plasmid |
| D70A_2-R | CAGATCCAGTTAATTGTGAG | To construct D68A+D70A and D68A+D70A+YVAA disruption plasmid |
| D70A-F | CTGGATCTGCTTTGACTTTGCTATAATTAAATCTAATTTAAATAG | To construct D70A point mutation plasmid |
| D70A-R | CTATTTAAATTAGATTTAATTATAGCAAAGTCAAAGCAGATCCAG | To construct D70A point mutation plasmid |
| En_G2-F | ACCATAGCATATGAAAATCGAAGAAGGTAAACTGGTAATC | To construct GIIxPPRT hybrid plasmid |
| En_G2-R | GCGCAACATTTCCTGCCCGCAGACTTTTC | To construct GIIxPPRT hybrid plasmid |
| En_PPRT-F | GCGGGCAGGAAATGTTGCGCTTAAAACATTAC | To construct GIIxPPRT hybrid plasmid |
| En_PPRT-R | CGATTTTCATATGCTATGGTCCTTGTTG | To construct GIIxPPRT hybrid plasmid |
| IntronPbF | CCTCTATCGACACATAACC | To construct targetron probe |

| | | |
|---|---|---|
| IntronPbR | CTACTCTGTAAGATAACACAG | To construct targetron probe |
| PPRT-DP1 | TCCAGCCATGGTTATACCTG | To amplify the down-stream of PRT |
| PPRT-DP2 | CTTGCCTGCAGTTAGTGATGATGATGATGATGATGATGCTCATTCAGTTCAAGCACTC | To amplify the down-stream of PRT |
| PPRT-UP1 | CGCCGCCATGGTTAACGAATTAGCCACATT | To amplify the up-stream of PRT |
| PPRT-UP2 | TATAACCATGGCTGGAATATG | To amplify the up-stream of PRT |
| PrmS-F | ATGGTGGATCCGTCGCAATC | To amplify Primase domain fragment |
| PrmS-R | GATTTGGCATGCTTAGACAGAG | To amplify Primase domain fragment |

**Table S6. Targetron probe**

| Sequence (5' to 3') |
|---|
| CTACTCTGTAAGATAACACAGAAAACAGCCAACCTAACCGAAAAGCGAAAGCTGATACGGGAACAGAGCACGGT TGGAAAGCGATGAGTTACCTAAAGACAATCGGGTACGACTGAGTCGCAATGTTAATCAGATATAAGGTATAAGTT GTGTTTACTGAACGCAAGTTTCTAATTTCGGTTATGTGTCGATAGAGG |
